# Language beyond the language system: dorsal visuospatial pathways support processing of demonstratives and spatial language during naturalistic fast fMRI

**DOI:** 10.1101/651257

**Authors:** Roberta Rocca, Kenny R. Coventry, Kristian Tylén, Marlene Staib, Torben E. Lund, Mikkel Wallentin

**Author notes:** Corresponding Author Email address Postal address: Jens Christian Skous Vej 2, Building 1485, Office 637, 8000 - Aarhus C, Denmark.

## Abstract

Spatial demonstratives are powerful linguistic tools used to establish joint attention. Identifying the meaning of semantically underspecified expressions like “this one” hinges on the integration of linguistic and visual cues, attentional orienting and pragmatic inference. This synergy between language and extralinguistic cognition is pivotal to language comprehension in general, but especially prominent in demonstratives.

In this study, we aimed to elucidate which neural architectures enable this intertwining between language and extralinguistic cognition using a naturalistic fMRI paradigm. In our experiment, 28 participants listened to a specially crafted dialogical narrative with a controlled number of spatial demonstratives. A fast multiband-EPI acquisition sequence (TR = 388ms) combined with finite impulse response (FIR) modelling of the hemodynamic response was used to capture signal changes at word-level resolution.

We found that spatial demonstratives bilaterally engage a network of parietal areas, including the supramarginal gyrus, the angular gyrus, and precuneus, implicated in information integration and visuospatial processing. Moreover, demonstratives recruit frontal regions, including the right FEF, implicated in attentional orienting and reference frames shifts. Finally, using multivariate similarity analyses, we provide evidence for a general involvement of the dorsal (“where”) stream in the processing of spatial expressions, as opposed to ventral pathways encoding object semantics.

Overall, our results suggest that language processing relies on a distributed architecture, recruiting neural resources for perception, attention, and extra-linguistic aspects of cognition in a dynamic and context-dependent fashion.

## 1. Introduction

### 1.1 Demonstratives: an interface between language, attention, and spatial cognition

The two utterances “I would like to buy the yellow cake” and “This one” can mean the same thing, depending on the circumstances. The latter is often used in situations where knowledge about the intended interaction (e.g. a buying frame) can be taken for granted, and the speaker simply wishes to point the hearer’s attention to the relevant object. Both sentences, however, use linguistic cues to coordinate interlocutors’ focus of attention to particular aspects of the environment. This ostensive function is a cornerstone of language, that supports collaboration and other forms of collective engagement with the physical world (Tomasello, Carpenter, Call, Behne, & Moll, 2005; Tylén, Weed, Wallentin, Roepstorff, & Frith, 2010).

Word types vary, as exemplified above, in the amount of semantic and extralinguistic (e.g. visuospatial) information needed for their comprehension. So-called *content* words, a category which includes most nouns, verbs, and adjectives, are expressions that denote objects (“cake”), qualities (“yellow”), or actions (“to buy”), by explicitly naming them. These expressions provide the semantic core of an utterance, as they have rich and view-point independent meaning (Diessel, 2006). Little extralinguistic information is needed to disambiguate the intended referent in the environment.

Other types of linguistic utterances, on the other hand, point to specific referents in the physical or discursive environment in specific situations, as seen from a specific viewpoint, without providing explicit semantic information about them. An example of this is spatial demonstratives, i.e. words like *this* and *that* in English. Demonstratives are *deictic* expressions (Levinson, 1983): when presented in isolation, they can denote virtually *any* referent. Interpreting what “this one” means hinges on perceptual processing (e.g. how far away from the speaker are potential referents located?), attentional orienting on the basis of gaze cues and pointing gestures (Cooperrider, 2016; García, Ehlers, & Tylén, 2017; Stevens & Zhang, 2013), and pragmatic inference (what could the speaker be intending to refer to?). Demonstratives are therefore a paradigmatic example of how linking language to the physical world requires the integration of linguistic forms with extra-linguistic perceptual and cognitive processing.

Demonstratives are foundational to language on a number of levels. They are linguistic universals (Diessel, 2014), they are milestones in language acquisition (Clark & Sengul, 1978), they are among the most frequent words in the lexicon (Leech & Rayson, 2014), and they play a crucial role in the evolution of grammar (Diessel, 2013). In spite of their importance in language, no neuroimaging studies investigating the neural processing of demonstratives exist, probably due to the methodological challenges posed by studying these words. As the meaning of demonstratives is dependent on the context, investigating their neural underpinnings hinges on simulating a rich linguistic and physical environment within the constraints intrinsic to neuroimaging experiments.

In this study, we constructed a novel naturalistic paradigm where we simulated such rich contexts, with the aim of elucidating which neural architectures enable the interaction between linguistic, perceptual, and attentional processes in language comprehension.

### 1.2 Usage patterns for demonstratives reflect functional encoding of space

The tight interdependencies between demonstrative reference and fundamental aspects of attention, perception, and spatial representations are explicitly reflected in usage patterns of different demonstrative forms.

The vast majority of natural languages encodes at least a binary distinction between a so-called *proximal* demonstrative, such as *this*, and a *distal* demonstrative form, such as *that* in English (Diessel, 1999). Experimental evidence has shown that this distinction does not encode purely metric distance between the speaker and the referent. In a series of experiments based on the *memory game* paradigm, Coventry and colleagues have shown that the contrast between proximal and distal demonstratives maps onto the functional distinction between *peripersonal* and *extrapersonal* space, that is, between space within and outside reach (Caldano & Coventry, in press; Coventry, Griffiths, & Hamilton, 2014; Coventry, Valdés, Castillo, & Guijarro-Fuentes, 2008; Gudde, Coventry, & Engelhardt, 2016). In these experiments, participants were presented with shapes placed at one of 12 potential distances along a table. After a (variable) number of trials, participants were presented again with one of the shapes, and asked to indicate at which location it had previously been placed by pointing at the intended location while producing phrases consisting of a demonstrative adjective, a color adjective and a noun (e.g. “this blue square”). Referents placed *within* reach were more likely to be indicated using proximal demonstratives, while objects *outside* reach were more likely to elicit distal demonstratives.

However, when multiple competing referents are present, their relative distance also matters when speakers choose between proximal and distal demonstratives. Bonfiglioli and colleagues conducted an experiment (Bonfiglioli, Finocchiaro, Gesierich, Rositani, & Vescovi, 2009) where participants were primed with either a proximal or a distal demonstrative forms before performing reach movements towards objects positioned at either of two possible distances *within* their peripersonal space. Semantic interference effects on movement initiation where detected (in terms of slower reaction times) when participants had to reach for the *closer* target location after being primed with a *distal* demonstrative, and when they reached for the *farther* location after being primed with a *proximal* demonstrative. The proximal/distal contrast thus also codes for relative distance between potential referents and the speaker even when *both* referents are within her reach.

Previous studies have furthermore detected lateralized biases towards the pointing hand. Speakers are more likely to use *proximal* demonstratives for referents located towards their right when pointing with their right hand, which has been interpreted as evidence in favor of a connection between demonstratives and affordances for manual action (Rocca, Wallentin, Vesper, & Tylén, 2018). A strong link between demonstratives and manual action has also been observed at a purely semantic level. When participants are asked to choose between a proximal and a distal demonstrative in the absence of any explicit spatial context, they consistently choose proximal demonstratives for objects that more easily afford manual grasp, such as small vs. big objects, and harmless vs. harmful referents (Rocca, Tylén, & Wallentin, 2019).

Additionally, demonstrative use is significantly modulated by social factors, such as the presence, position, and role of an interlocutor in the ongoing interaction. On the basis of the results from an EEG/ERPs study, Peeters and colleagues have argued that the use of proximal vs. distal demonstrative forms is influenced by whether the referent is located within vs. outside the region of space shared between two interlocutors (Peeters, Hagoort, & Özyürek, 2015). Other studies have found that speakers tend to adapt their use of demonstratives to the position of the addressee in the context of social interaction. During collaborative interactions, proximal space tends to shift towards the addressee (Rocca et al., 2018), Speakers tend to code locations as proximal or distal depending on their distance from the *addressee*, rather than from themselves, in interactions involving turn-taking (Rocca, Wallentin, Vesper, & Tylén, 2019).

In summary, behavioral evidence suggests that the use of demonstrative forms is influenced by extralinguistic perceptual, functional and social representations of space. This leads us to hypothesize that similar extralinguistic representations might be necessary on the addressee’s side, in order to process the cues provided by the use of proximal vs. distal forms.

### 1.3 A dorsal pathway for semantics?

Previous literature on spatial language has suggested that processing spatial expressions shares resources with non-linguistic spatial encoding. A network of dorso-parietal brain regions supports both visuospatial perception and linguistic reference to the perceived space (Wallentin, Roepstorff, Glover, & Burgess, 2006; Wallentin, Weed, Østergaard, Mouridsen, & Roepstorff, 2008), while shifting spatial frames of references engage the system for shifting visual attention, including the frontal eye fields (Corbetta et al., 1998; Wallentin, Kristensen, Olsen, & Nielsen, 2011; Wallentin, Roepstorff, & Burgess, 2008). Additionally, integration areas in the inferior part of the parietal lobe, namely the left supramarginal gyrus and the angular gyri, have been implicated in processing of spatial closed class items, such as prepositions (H. Damasio et al., 2001; Kemmerer, 1999, 2006; Noordzij, Neggers, Ramsey, & Postma, 2008). The SMG is part of the temporoparietal junction, which interfaces the auditory cortex with parietal and frontal circuits (Scott & Johnsrude, 2003). The angular gyrus, located in the IPL, has been implicated in complex information integration and knowledge retrieval (Binder, Desai, Graves, & Conant, 2009) and in scene construction (Hassabis & Maguire, 2009), both central in processing linguistic spatial relations.

Interestingly, posterior-superior parietal areas and frontal regions identified in previous studies on spatial language all belong to the dorsal visuo-spatial stream (Mishkin, Ungerleider, & Macko, 1983). This suggests that, globally, language processing might be organized along a ventral-dorsal divide between semantics and (spatial) relations parallel to that between object identification and locations in vision (Landau & Jackendoff, 1993, 2013). Naming objects and talking about their locations differ widely in the type of information encoded in linguistic forms. Object descriptions draw on abstract representations of spatial features, prioritizing viewpoint-independent attributes such as shape and surface relevant to categorization. Spatial relations, on the other hand, are conveyed by very coarse geometrical detail, mostly drawing on functional properties such as relative distance, containment, and contact. This provides sufficient cues for allocating attention to the relevant part of space or time in order to access more detailed information.

The hypothesis of a ventral/dorsal what/where divide in language is supported by evidence from semantic analyses of linguistic expressions and the studies mentioned above, but whether such a divide is rooted on a functional segregation at the neural level has never *directly* been tested empirically. In our study, we aimed not only to elucidate the neural architecture underlying the processing of spatial demonstratives, but also at directly testing the hypothesis of the existence of a dorsal “where” stream for the processing of linguistic spatial relations, largely overlapping with the visuospatial dorsal stream.

Such results would make a compelling empirical case in favor of a ventral-dorsal segregation in language processing, and, more generally, underline the what/where distinction being a fundamental organizational principle for information processing in the human brain.

### 1.4 Present study: experimental paradigm

In this experiment, we presented participants with a specially crafted, scripted dialogue featuring two voices (a male and a female). The decision to use dialogue was motivated by the fact that, as demonstratives are prominently used to establish joint attention, they tend to occur in dialogic contexts, rather than in monologues or written discourse (i.e. *that* is 5.5 times more frequent in spoken language than in written, and *this* is 1.2 times more frequent, see Leech & Rayson, 2014). The choice of spoken dialogue therefore added further ecological validity to our investigation.

In the dialogue, two characters try to find each other in the darkness, a setting which naturally affords occurrences of spatial expressions. Demonstratives can be used *exophorically*, i.e. to refer to objects in the perceptual environment, or *endophorically*, that is, in an intralinguistic fashion, to denote parts of discourse (Diessel, 1999). This study focuses on the *exophoric* use.

Several demonstratives were inserted in the text, with a balanced number of proximal (*here*) and distal (*there*) demonstratives, equally distributed across voices. By recording the two voices onto two separate audio channels, we simulated a minimal 3D-like auditory environment where participants experienced one character as being located to their left and the other to their right. Demonstratives provide indications on the position of objects (or locations) relative to the position of the speaker and conversational dyad (Coventry et al., 2014, 2008; Gudde et al., 2016; Peeters et al., 2015). It is therefore crucial that the two speakers in the dialogue are assigned specific and distinct spatial origins.

Moreover, this manipulation enabled us to tease apart the effect of different demonstrative forms (here vs. there) from the effects of the location they denote in auditory space (left, right), especially with regards to proximal demonstratives. The location denoted by proximal demonstratives is tied to the position of the speaker and interacts with the spatial source of the speech input (while the scope of distal demonstratives is broader).

Our paradigm relied on a fast acquisition sequence (TR = 388ms), which, combined with finite impulse response (FIR) modelling of the hemodynamic response, allows us to optimally capture neural response at word-level resolution within naturalistic paradigms even when response patterns deviate from the time course of the canonical hemodynamic response function. Deviations from the canonical response model might indeed be expected on the basis of recent results showing that, under sustained stimulation (of which naturalistic speech is an instance), the hemodynamic response is faster than assumed by the canonical HRF (Lewis, Setsompop, Rosen, & Polimeni, 2016).

### 1.5 Hypotheses

In our analysis, we tested the following hypotheses:

First, we investigated which brain areas respond to the occurrence of spatial demonstratives, averaging across proximal and distal demonstrative forms. We hypothesized that processing spatial demonstratives would engage a) areas interfacing the speech input with visuospatial processing in the parietal lobes, such as the supramarginal gyrus (Scott & Johnsrude, 2003); b) higher-order integration areas in the posterior parietal cortex such as the angular gyrus, previously implicated in tasks requiring complex information integration (Binder et al., 2009; Hassabis & Maguire, 2009) and therefore likely crucial for spatial demonstratives, where comprehension hinges on integrating the categorical distance cues with the visuospatial, linguistic and pragmatic context. The left SMG and AG have been previously implicated in the processing of spatial prepositions (H. Damasio et al., 2001; Kemmerer, 1999, 2006; Noordzij et al., 2008). Moreover, we expected demonstratives to engage c) medial parts of the superior posterior parietal cortex, previously implicated in constructing and maintaining spatial representations for both language and vision (Wallentin et al., 2006; Wallentin, Weed, et al., 2008), and d) frontal regions within the dorsal parieto-frontal attentional network effecting the attentional shifts triggered by spatial demonstratives (Corbetta et al., 1998; Wallentin et al., 2011; Wallentin, Roepstorff, et al., 2008).

Second, we compared proximal and distal demonstratives, exploring differences in the neural correlates of the two forms. Behavioral evidence on demonstratives suggests a mapping between demonstrative forms and the distinction between peripersonal and extrapersonal space. Differences between proximal and distal forms might therefore be encoded in the superior parietal lobule (SPL) and superior parieto-occipital cortex (SPOC), previously implicated in spatial encoding for manual reach (Andersen, Andersen, Hwang, & Hauschild, 2014; Connolly, Andersen, & Goodale, 2003; Gallivan, Cavina-Pratesi, & Culham, 2009; Grivaz, Blanke, & Serino, 2017).

Additionally, we analyzed interactions between demonstrative form and ear of presentation. In line with preferences for contralateral locations observed in the frontoparietal attentional stream (Halligan, Fink, Marshall, & Vallar, 2003), we tested whether areas responding to demonstratives displayed higher sensitivity to proximal forms in the contralateral ear and distal forms in the ipsilateral ear, i.e. to cases where demonstratives likely code for locations in the contralateral spatial hemifield.

Third, we tested whether, more generally, neural processing of spatial relations (as expressed in language) relies on a dorsal *where* processing stream, as opposed to a ventral *what* stream for object semantics. To test this hypothesis, we compared response to spatial demonstratives with response with the wh-words *where*, *what*, and *who*. These words prime the processing of spatial information, object identity, and personal identity respectively, and therefore function as proxies to the divide between semantic content and spatial relations in language. Neural representations for these words were compared to representations underlying demonstratives using a novel similarity-based method, under the hypothesis of higher topographical similarity between demonstratives and *where* at the whole-brain level. Zooming in on an anatomical partitioning of brain areas, we expected this pattern to be mostly driven by higher topographical similarity in areas belonging to the dorsal processing stream. If this hypothesis held true, this would suggest that resources supporting language processing strongly overlap with resources for visuo-spatial processing, inheriting fundamental organizational principles (dorsal vs. ventral) shared across multiple domains of human cognition.

Besides testing these hypotheses, we ensured that our acquisition sequence yielded high-quality images by regressing the data against low-level acoustic features (sound envelopes from both audio channels), expecting to replicate results from previous literature (Jäncke, Wüstenberg, Schulze, & Heinze, 2002; Schönwiesner, Krumbholz, Rübsamen, Fink, & von Cramon, 2006; Stefanatos, Joe, Aguirre, Detre, & Wetmore, 2008) on spatial activation patterns in the auditory cortices for monaural stimulation. We expected both auditory cortices to respond to both envelopes for the left and right auditory channels, with larger and more widespread response in the contralateral auditory cortex. Additionally, exploiting the combination of high sampling rate (∼2.58Hz) with flexible FIR models, we explored temporal BOLD response patterns in auditory cortices under sustained speech stimulation.

## 2. Methods

### 2.1 Participants

Twenty-nine participants with normal hearing and anatomically normal brains took part in the study. Data from one participant were discarded from the analysis, due to the presence of artifacts in the EPI images. Therefore, data from 28 participants (Female = 12, Age median = 24, Range = 19-36) were included in the analyses. Participants were recruited on a voluntary basis from the participant pool of the Center for Functionally Integrative Neuroscience at Aarhus University. All participants were right-handed and reported having Danish as their first language. Gender was not deemed relevant (Wallentin, 2009, 2018). The study received approval from the research ethics committee of Region Midtjylland, Denmark, and participants gave informed written consent in accordance with local ethical requirements. Participants received monetary compensation for their participation in accordance with local policies on participant payment.

### 2.2 Acquisition details

Functional images were acquired on a 3-T Siemens Magnetom Tim Trio MR system equipped with a 32-channels head coil at Aarhus University Hospital, Denmark. For each participant, 3670 volumes, each containing 54 T2*-weighted slices, were acquired using a multiband-EPI sequence, with repetition time (TR) = 388ms, echo time (TE) = 27.6ms, flip angle: 36°, voxel size = 2.5mm isotropic, slice-acceleration factor = 9 (Setsompop et al., 2012), but no in-plane acceleration.

At the end of each session, a gradient echo-based field map was acquired, based on subtraction of the phase images from a dual echo acquisition, with the following parameters: repetition time (TR) = 1020ms, echo time (TE) = 10ms and 12.46ms, flip angle = 90°, voxel size = 3mm isotropic, field of view =192 x 192 mm. These field maps were then used to unwarp geometrical distortions due to field inhomogeneities using the FieldMap toolbox and the Unwarp module in SPM12.

Pulse-oximetry and respiration were recorded during the whole experiment using scanner hardware, and used for denoising purposes. Modelling cardiac and respiration data in GLM analyses has proven effective in accounting for serial correlations in the noise structure of EPI time series, especially in the context of acquisition sequences with sub-second temporal resolution (Bollmann, Puckett, Cunnington, & Barth, 2018; Lund, Madsen, Sidaros, Luo, & Nichols, 2006; Purdon & Weisskoff, 1998; Sahib et al., 2016).

### 2.3 Stimuli

Participants listened to a spoken dialogue (in Danish) with a total duration of 23 minutes and 40 seconds through headphones. No visual stimuli were displayed during the experiment. Participants were instructed to keep their eyes open through the experiment.

In the dialogue, two fictive characters are heard, one speaking through the left channel of the headphones and the other speaking through the right. The two characters find themselves in a dark and unfamiliar environment. The dialogue unfolds with constantly alternating focus on narrative and spatial information. Over the course of the interaction, the two characters try to figure out where they are, what the surrounding environment looks like, who their interlocutor is, as well as how and why they ended up in the darkness. This setting, where characters are constantly engaged in exploring and describing a spatial scene, makes room for several motivated occurrences of spatial demonstratives. Moreover, it provides a suitable context for questions, and therefore wh-words, to occur naturally and with high frequency.

These characteristics enabled us to a create naturalistic speech stimulus while retaining control of the frequency of occurrence of words of interest, as well as on their position and spacing in the text.

The full text of the dialogue in Danish and in an English translation is available at <u>osf.io/j9fm5/</u>. Overall, the dialogue included 80 occurrences of each demonstrative form (*proximal* = *her*, *distal* = *der*), equally distributed across the two voices (and therefore auditory hemifields). Inter-stimulus intervals for each demonstrative type were not fixed but semi-controlled, with a mean ISI of 17.78s for proximal demonstratives and a mean ISI of 17.43s for distal demonstratives. Forty instances of the words *what* (*hvad*), *where* (*hvor*), and *who* (*hvem*) were embedded in the text, balanced across the two voices. The mean ISI was 31.39s for *what,* 35.76s for *where*, and 33.7s for *who*.

The dialogue unfolds over 340 lines (170 per character). The two characters speak a total of 1585 words and 1470 words.

One hundred instances of singular first- and second-person pronouns (*I* and *you*) also occurred in the text, equally distributed across voices. The results of this latter manipulation will be reported elsewhere.

### 2.4 Speech synthesis

The dialogue was recorded using two synthesized Danish voices (a male and a female). We interfaced an NSSpeechSynthesizer instance on macOS Sierra (Version 10.12.2) via the *pyttsx* library. The script set each voice to read aloud specific parts of the dialogue at a pace of 130 words per minute. The sound output was played and recorded on the internal audio system using SoundflowerBed (v 2.0.0) and saved as waveform stereo file with a sampling rate of 44.1kHz. We embedded AppleScript commands interacting with QuickTime Player (v 10.4) in the Python script, in order to automatize recording and time-lock the audio file to the onset of the sound stimulus.

Using text-to-speech synthesis offered a number of advantages over using recordings of natural voices. The engine interface in *pyttsx* allowed us to implement a callback function providing exact time stamps for the onset of each word in the dialogue. This overcomes the disadvantages of manual coding of audio files both in terms of precision and time requirements. Moreover, speech synthesis enabled an optimal combination of control and flexibility in stimulus generation. The output was tightly controlled in terms of pace and pronunciation, and the audio signal was not affected by any source of noise. Overall, the automatization of stimulus generation using Python-based speech synthesis enabled us to flexibly refine our stimulus over different steps of the piloting process, optimizing time demands over repeated iterations of processing and annotation stages.

The dialogue was recorded onto a two-channel stereo track, with each voice presented monaurally. Manipulating the spatial source of voices afforded simulation of a minimal 3D spatial context, with each character being experienced as located either to the left or to the right of the participant.

The dialogue was presented through MR-compatible OptoACTIVE headphones (OptoAcoustics Ltd.). The side of presentation of each voice was counterbalanced across participants.

### 2.5 Online behavioural task

During the experiment, participants performed a simple on-line behavioural task, to ensure that they remained actively engaged throughout the experiment and to avoid data loss due to participants falling asleep. Thirty breaks lasting 5 seconds were embedded in the dialogue. Fifteen out of thirty breaks were interrupted by a pure tone of 500ms duration. Participants were instructed to respond to the occurrence of pure tones by pressing a button on the response box.

Tones always occurred during silent breaks, and their onset followed the start of the break with a perceptible lag. Participants were informed that tones would only occur during the silent breaks, so to make sure that they could entirely focus on the comprehension of the dialogue without expecting sudden disruptions of its flow. Participants were split into two groups. Groups differed in the subset of breaks during which pure tones were presented in order to decorrelate perceptual and motor effects from the linguistic stimuli across participants. PsychoPy2 (Peirce, 2007) was used for stimulus delivery and response collection.

Twenty-six (26) out of 28 participants responded to all tones embedded in the dialogue, while the remaining 2 participants responded to 14 out of 15 tones. Performance levels for all participants were therefore deemed sufficient for inclusion in the analysis.

### 2.6 Post-experiment behavioural tasks

Participants performed two additional post-experiment tasks outside the scanner. Before entering the scanner, participants were informed that, at the end of the experiment, they would be asked to draw the scene where the dialogue took place, and answer some comprehension questions on the content of the dialogue. While responding to tones ensured general engagement during the unfolding of the experiment, the post-experiment tasks motivated participants to pay close attention to the content of the dialogue and tested their actual comprehension of the text.

The drawing task was meant to prime participants to focus on spatial expressions, while still keeping them naïve to our interest in spatial demonstratives. Drawings were entirely unconstrained in terms of degree of detail, number of elements represented, and their configuration. No behavioural metrics were extracted from this task.

The questionnaire tested engagement in the comprehension of the dialogue, and it was meant to provide a behavioural criterion for inclusion in the fMRI analysis. Participants answered 20 comprehension questions tapping onto narrative aspects of the stimulus story, e.g. information on characters and events mentioned during the dialogue. All participants performed significantly above chance (mean performance = 88.2% correct responses) and were therefore included in the fMRI analysis.

After the comprehension questionnaire, participants were administered a short questionnaire tapping onto their experience of the dialogue. All participants reported being able to hear the two voices clearly and to understand the dialogue without major effort. They were explicitly asked to comment on whether and how the use of synthesized voices affected their experience of the dialogue. Some participants reported having noticed a few oddities in the pronunciation, but all specified that this did not have an impact on their comprehension of and focus on the content. No participants reported tones being disruptive of their engagement in the comprehension of the dialogue.

### 2.7 Data pre-processing

#### 2.7.1 EPI images and anatomical images

Data were preprocessed using SPM12. T1-weighted images, T2*-weighted EPI images and field maps were first converted from DICOM to NIFTI format. EPI images were then realigned to the first image in the time series via rigid body spatial transformations. Realignment parameters for each subject were stored and used in the GLM analyses to account for residual movement-related variance.

Using the FieldMap toolbox, subject-specific voxel displacement maps were computed from the presubtracted phase image and the magnitude image with shorter echo time (TE = 10ms). EPI images were then unwarped using the resulting voxel displacement maps to correct for geometric distortions caused by field inhomogeneities. Subject-specific anatomical images were co-registered to the mean unwarped functional image, then segmented into 6 tissue maps. A 4mm FWHM smoothing filter was applied to the images prior to estimation of a forward deformation field, used to normalize the unwarped EPI images and T1-weighted images to MNI space.

#### 2.7.2 Physiological data

Pulse-oximetry and respiration data were processed using Matlab PhysIO Toolbox (Kasper et al., 2017) and modelled using the RETROICOR algorithm (Chang, Cunningham, & Glover, 2009; Glover, Li, & Ress, 2000) with 3^rd^ order and 4^th^ order expansion for cardiac and respiratory terms, and 1^st^ order expansion for their interaction. The 6 movement regressors estimated during realignment of EPI images were included in the RETROICOR model, and all regressors were orthogonalized.

### 2.8 Hemodynamic response modelling

In all GLM analyses reported in the Results section, hemodynamic response was modelled using finite impulse response (FIR) basis sets including 20 basis functions with 20 contiguous 500ms time bins modelling hemodynamic response from 0 to 10 seconds after stimulus onset.

FIR basis sets model the average peristimulus signal over each time bin via linear deconvolution of impulse response (Henson, 2003). Carrying minimal assumptions on the response, FIR models allow for local variation in its shape and amplitude, and can capture event-related signal changes with temporal patterns that deviate from the canonical HRF. Coupled with fast acquisition protocols, FIR models thus enable detection of high-frequency modulations present in the BOLD signal under sustained fast-paced stimulation (Lewis et al., 2016). This makes these models suitable for naturalistic experiments on word semantics, where the speech rate of the stimulus tends to exceed one hundred words per minute, and responses to individual lexical units are likely expressed by high-frequency modulations over a sustained response.

### 2.9 GLM analyses

#### 2.9.1 Model structure and statistical inference

In all GLM analyses reported in the Results section, first-level models included regressors coding for the occurrence of each event of interest (differing across analyses), and a shared set of regressors accounting for non-speech events occurring in the experiment (silent breaks, pure tones, button presses). All components of individual RETROICOR models for physiological data and the 6 realignment parameters were added as covariates to account for residual movement-related variance and physiological noise.

For all analyses, T-contrasts testing for the effects of interest were computed on the first level, and contrast images at each time bin were entered into a second-level ANOVA with non-sphericity correction. The second-level model included 28 contrast images (one per subject) for each time bin, as well as covariates accounting for subject-specific effects.

Group-level inference was based on F-contrasts testing for the significance of first-level estimates at any time bin. The results of these contrasts were masked so to include only those voxels which are also significant in T-contrasts testing for an average positive effect across the 10 seconds post-stimulus interval. This allowed us to limit inference to those regions where signal increased as a response to events of interest, as well as to exclude those regions where F-tests might capture unreliable effects driven by estimates in one (or few) time bins. Such masking was not applied when directly testing for differences between regressors, thus allowing for effects with both positive and negative directionality.

In all analyses, inference was drawn at the voxel level using a significance threshold of p<.05 (FWE-corrected) and an additional spatial extent threshold of 30 voxels.

#### 2.9.2 Sound envelope

In order to ensure that the fast acquisition sequence yielded high-quality EPI images, we performed a first whole-brain analysis targeting responses to low-level acoustic features (sound levels) of the stimulus, expecting significant effects in the auditory cortices, with larger and more widespread response in the contralateral auditory cortex (Jäncke et al., 2002; Schönwiesner et al., 2006; Stefanatos et al., 2008). In this analysis, sound envelopes for the left and the right channels were used as regressors of interest in the first-level models.

#### 2.9.3 Spatial demonstratives

In the GLM analyses testing for regions responding to the occurrence of spatial demonstratives, four sets of FIR regressors were included at the first-level, modelling the onsets of *proximal* and *distal* demonstratives in the *left* and the *right* auditory channel. The model thus included one set of FIR predictors coding for *all* occurrences of *proximal demonstratives* in the *left* auditory channel, one set coding for *all* occurrences of *distal demonstratives* in the *right* auditory channel, one set coding for *all* occurrences of *proximal demonstratives* in the *right* auditory channel, one set coding for *all* occurrences of *distal demonstratives* in the *right* auditory channel. As all non-speech events in the experiment (nuisance regressors, silent breaks, tones, button presses) were also modelled, what was left unmodelled was thus a general language baseline. Analyses testing for an average effect of demonstratives thus compare demonstratives to this general language baseline.

#### 2.9.4 Wh-words

To extract parameter maps for wh-words for multivariate similarity analyses, we fitted a comprehensive model including regressors for all words systematically manipulated in the experiment. This model included two sets of FIR regressors coding for all occurrences of proximal and distal demonstratives (*here* and *there*, regardless of side of presentation), three sets coding for all occurrences of wh-words (*what*, *where*, *who*) and regressors coding for occurrences of personal pronouns. First-level parameter estimates for each demonstrative and wh-word were used as input to compute correlations used in the multivariate similarity analysis.

### 2.10 Multivariate similarity analyses

First-level FIR models yielded, for each regressor and for each participant, one parameter map for each post-stimulus time bin. From the cumulative model including regressors for all experimentally controlled words, we extracted parameter maps for the two demonstrative forms (*proximal* and *distal*), and for the wh-words *where*, *what* and *who*, at each time point. This yielded 28 (subjects) x 20 (time points) x 5 (words) parameter maps.

For each subject and at each of the 20 time points, we computed Pearson’s correlations between parameter maps for demonstrative forms and wh-words. Correlations between whole-brain parameter maps for each pair of words quantified global topographical similarity in response to such words at each time point. This yielded one correlation value for each of the 28 subjects, at each of the 20 time points, for each of the 6 combinations between a demonstrative and a wh-word.

As expanded upon in the Results section, three summary metrics (area under the curve, mean correlation, and maximum correlation) were extracted for each correlation time series. These measures were used as outcome variables in linear mixed-effects regression models comparing whole-brain topographical similarity between representations of demonstratives and representations of each wh-word. Zooming in on similarity patterns at a more local level, we also computed Pearson’s correlations for each word pair, each subject, and each time point on 60 brain regions extracted from the AAL atlas (Rolls, Joliot, & Tzourio-Mazoyer, 2015; Tzourio-Mazoyer et al., 2002; see also Appendix A for more details). A descriptive overview of local topographical similarity patterns is provided in the Results section.

### 2.11 Data and Code Availability Statement

Materials and code for the present experiment are publicly available on the Open Science Framework (<u>osf.io/j9fm5/</u>). The repository includes the full text of the stimulus dialogue in Danish and a full English translation, the audio files used as stimuli (in Danish), a 5 minutes audio sample in English, Python scripts used for stimulus creation and delivery, processed fMRI data and analysis scripts for both whole-brain and ROI-based similarity analysis, English translations of the post-experiment questionnaires, data and analysis script for the post-experiment comprehension questionnaires, data and analysis scripts for the online behavioural task. The repository also includes a description of each item in its wiki. Raw MR data are not fully anonymized and have therefore not been made publicly available.

All the group-level statistical maps (both thresholded and unthresholded) are publicly available on NeuroVault, at the ID: https://identifiers.org/neurovault.collection:5717.

Analysis scripts are available on GitHub at: <u>rbroc/demonstrativesfMRI</u>. GLM models for first-level analyses and second-level results can be shared upon request.

## 3. Results

### 3.1 Univariate analyses

#### 3.1.1 Sound envelopes

Variation in sound levels in the left channel significantly modulated activity in the right auditory cortex, with peak in the primary auditory cortex and extending along the superior temporal gyrus, MNI: [52, −18, 6], F_20,513_= 30.07, cluster extent = 1220 voxels, and in the left auditory cortex, peak MNI coordinates: [−66, −24, 0], F_20,513_= 15.65, cluster extent = 124 voxels.

Additional clusters in the precentral and postcentral gyri also responded to modulations in the left sound envelope. We detected significant clusters in the right precentral gyrus, peak MNI: [54, −8, 50], F_20,513_= 14.77, cluster extent = 43 voxels, and left precentral gyrus, peak MNI: [−54, −14, 50], F_20,513_= 8.02 and cluster extent = 46 voxels. Significant clusters were also detected in the right postcentral gyrus, peak MNI: [22, −38, 74], F_20,513_ = 12.20, cluster extent = 52 voxels, and left postcentral gyrus, peak MNI: [−58, −26, 46], F_20,513_= 8.00, cluster extent = 31 voxels.

**Figure 1.**
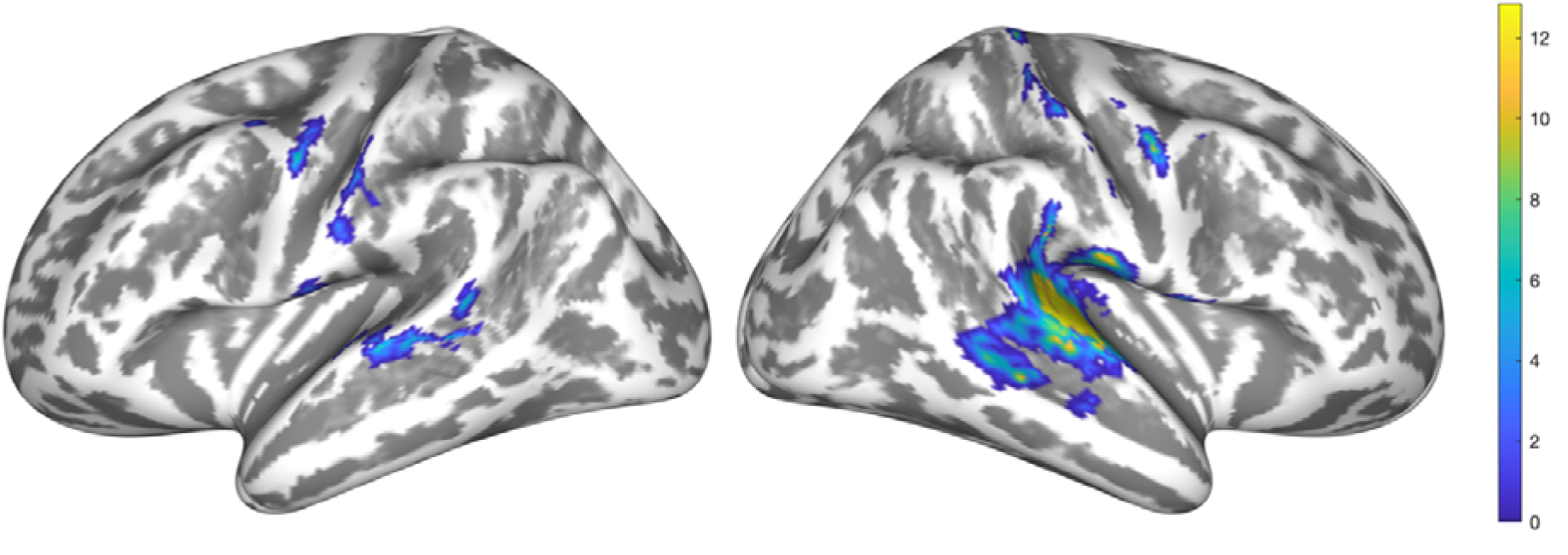
Regions responding to variation in sound levels in the left audio channel, p(FWE) < .05, cluster threshold = 30 voxels. Colors code for F-values (df = 20, 513) from second-level contrasts.

Figure 2 displays the time course of contrast estimates at each contiguous 500ms time bin after stimulus onset. The time course of the response is similar across hemisphere, with peak between 3 and 4 seconds, but the intensity of the response was higher in the contralateral auditory cortex.

**Figure 2.**
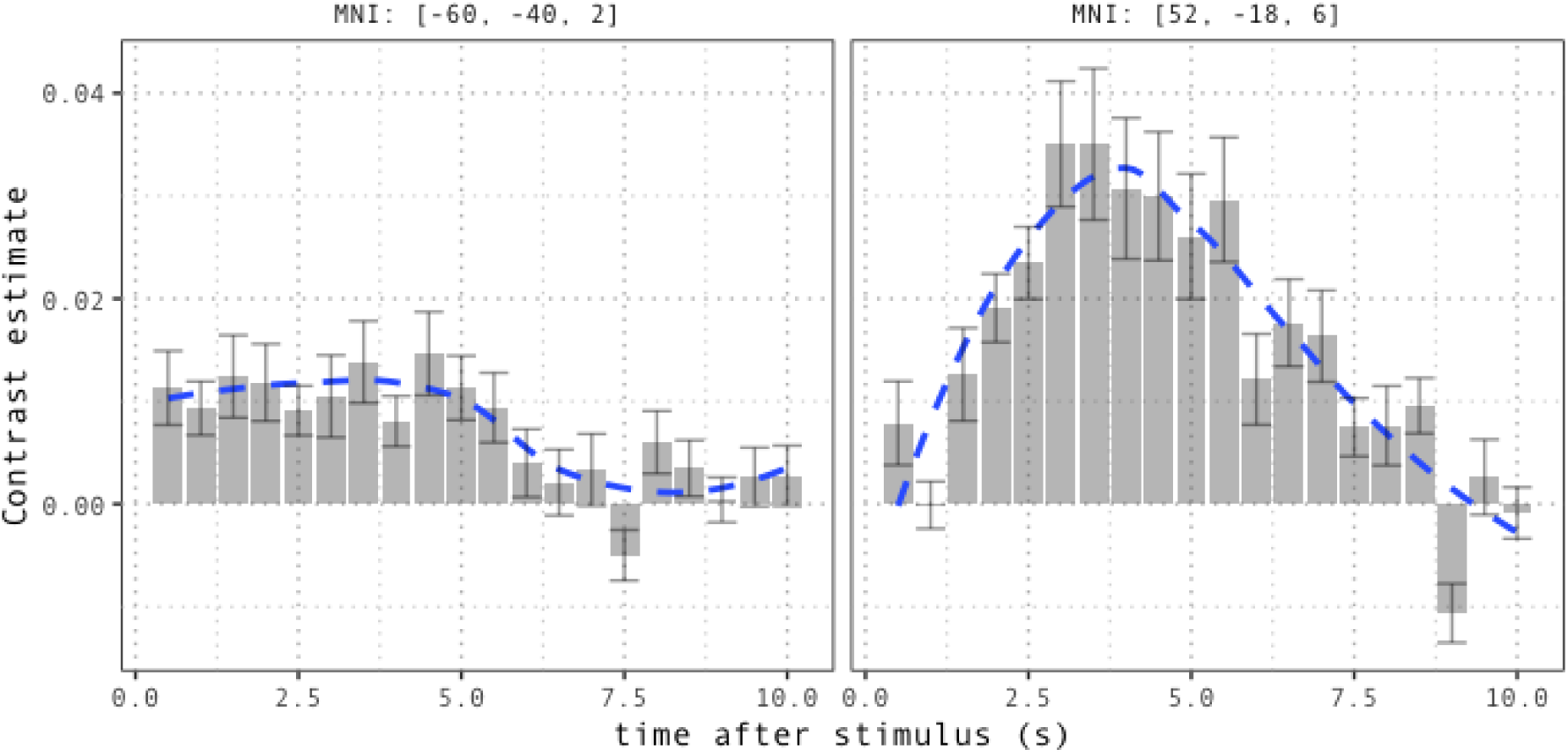
Time course of the response to variation in sound levels in the left channel at peak voxels in the left auditory cortex (left panel), and right auditory cortex (right panel). Error bars indicate between-participant standard errors. The overlaid curve smooths the average time series using local regression.

Sound levels in the right channel significantly modulated response in the left auditory cortex, with peak in A1 at MNI coordinates [−52, −28, 10], F_20,513_= 44.27, cluster extent = 1722 voxels, and in the right auditory cortex, peak MNI: [64, −12, 8], F_20,513_= 23.69, cluster extent = 898 voxels.

Beyond auditory cortices, we detected clusters with peaks in the left precentral gyrus, peak MNI coordinates [−52, −10, 48], F_20,513_= 24.94, cluster extent = 250 voxels, in the right precentral gyrus, peak MNI coordinates [56, 2, 44], F_20,513_= 14.62, cluster extent = 120 voxels, and in the left postcentral gyrus, peak MNI coordinates [−56, −26, 50], F_20,513_= 9.92, cluster extent = 42 voxels.

**Figure 3.**
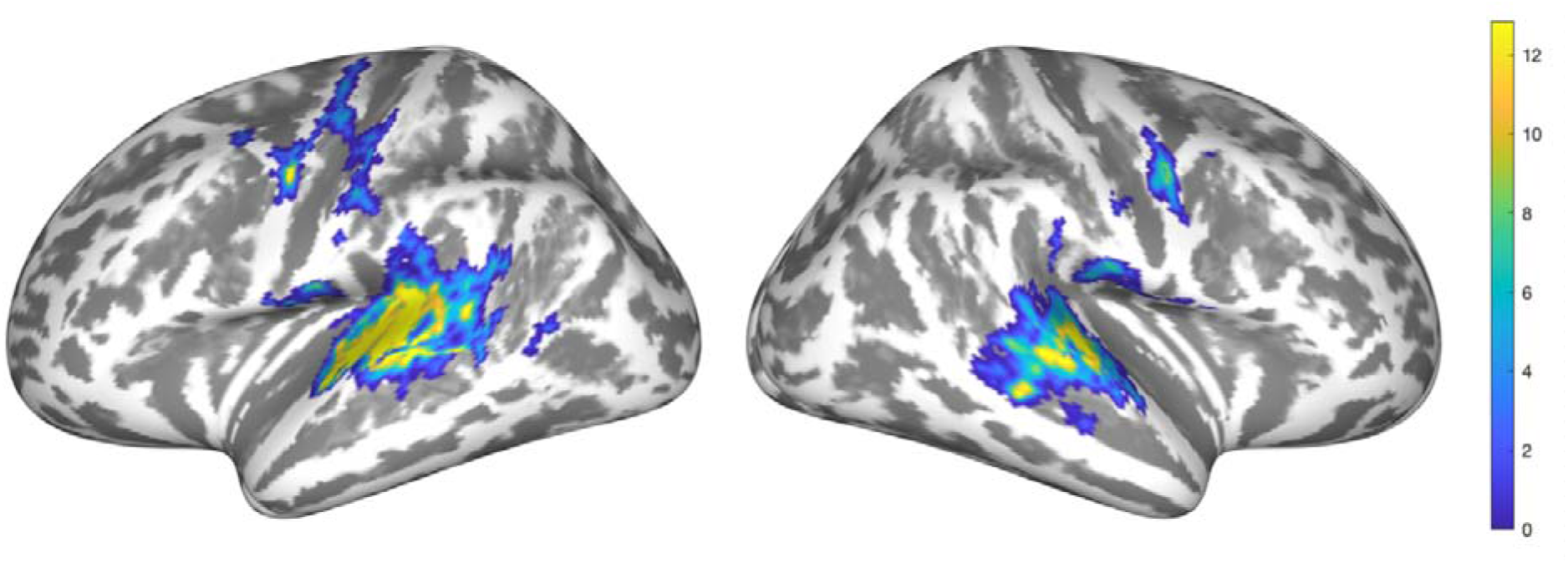
Regions responding to modulations in sound levels in the right channel. Significant clusters were detected in the left and right auditory cortices, as well as in the precentral gyri bilaterally and in the left postcentral gyrus. Colors code for F-values (df = 20, 513) from second-level contrasts.

As observed for the left sound envelope, response was larger in the contralateral auditory cortex. Contrast estimates show that response peaks around 3-4 seconds in the left auditory cortex, while response might peak later in the right auditory cortex.

**Figure 4.**
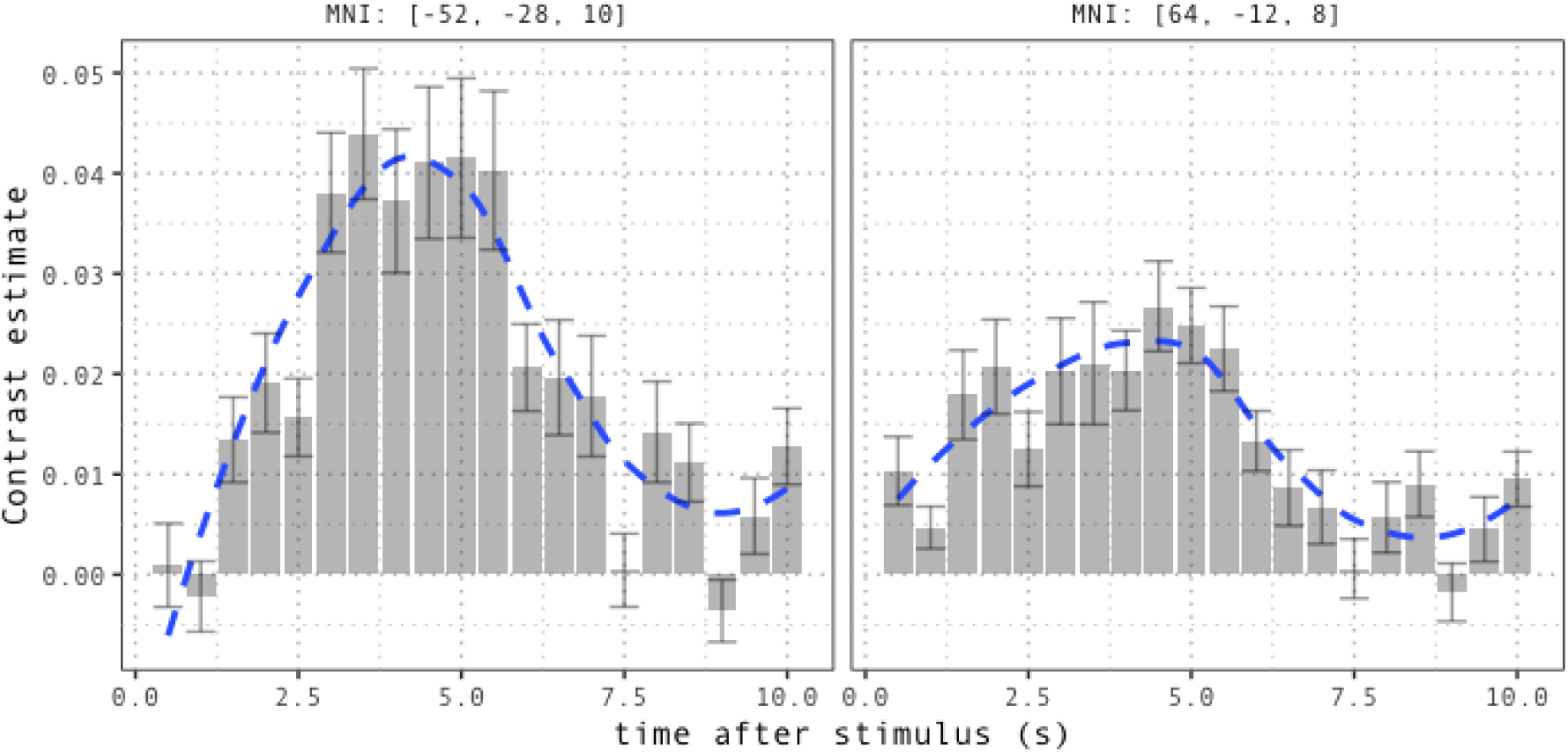
Time course of the response to variation in sound levels in the right channel, for peak voxels in the left auditory cortex (left panel), and right auditory cortex (right panel). Error bars indicate between-participant standard errors. The overlaid curve smooths the average time series using local regression.

A direct comparison of response to the left and right sound envelope showed that the left auditory cortex displayed a stronger response to variation in sound levels in the right channel. The contrast detects a cluster with peak in the left primary auditory cortex, MNI: [54, −28, 10], F_20,513_ = 26.42, cluster extent = 986 voxels. No preference for either the contralateral or the ipsilateral auditory hemifield was observed for the right auditory cortex.

Overall, the observed lateralization patterns, with bilateral responses marked by a contralateral advantage, are in line with our prediction. Moreover, our results suggest that, in the context of monaural stimulation, the magnitude of response to auditory stimuli in the right hemifield is stronger, which is consistent with the right-lateralized advantages in auditory processing largely attested in the literature (Hugdahl & Westerhausen, 2016; Kimura, 1967). These results are discussed in more detail in Appendix B.

**Figure 5.**
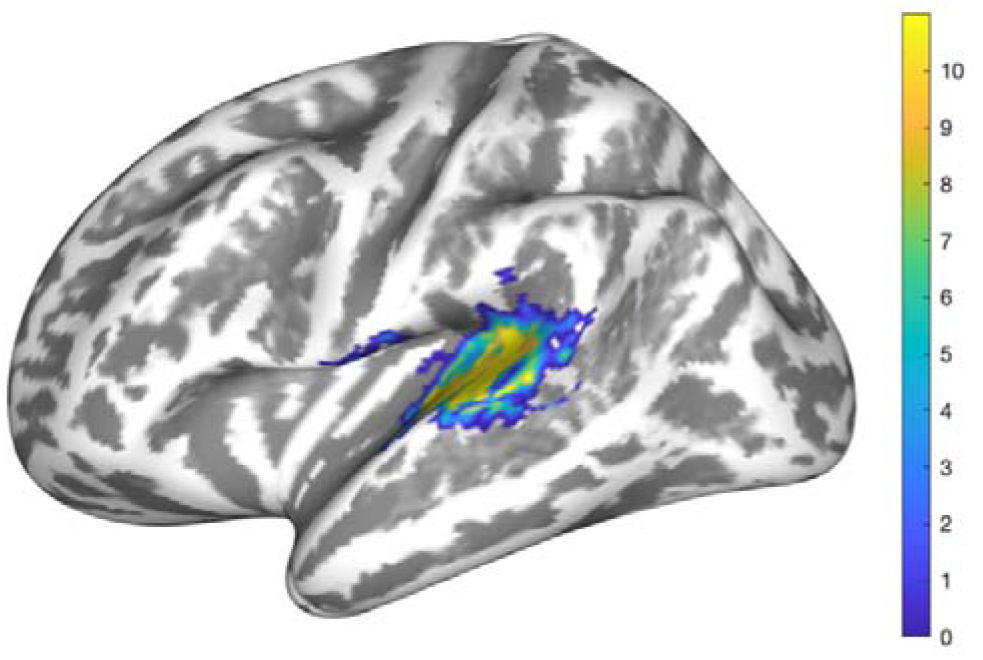
Direct contrast between left and right sound envelope. The left auditory cortex responds more strongly to sound variations in the right channel than in the left channel. No asymmetry is observed in the right auditory cortex.

#### 3.1.2 Demonstratives

##### 3.1.2.1 Average effect of demonstratives (proximal and distal)

The occurrence of demonstratives across both sides of presentations significantly modulated activity in a bilateral network involving inferior parietal, frontal and parieto-occipital regions.

In the inferior part of the parietal lobes, we detected a cluster with peak in the posterior part of the left angular gyrus, MNI: [−38, −80, 36], F_20,513_= 29.68, cluster extent = 362 voxels, and a cluster with peak in the right angular gyrus, MNI: [40, −74, 42], F_20,513_= 23.40, cluster extent = 439 voxels, both extending towards the middle occipital cortex. We also detected significant activation in the left supramarginal gyrus, peak MNI coordinates [−42, −50, 58], F_20,513_= 12.00, cluster extent = 67 voxels.

Demonstratives also modulate activity in the left precuneus, peak MNI coordinates [−2, −78, 42], F_20,513_= 11.77, cluster extent = 131 voxels, and in the right precuneus, peak MNI coordinates [10, −76, 42], F_20,513_= 9.71, cluster extent = 34 voxels.

The anterior part of the middle frontal gyrus also responds to the occurrence of demonstratives, with a significant cluster in the left hemisphere, peak MNI coordinates [−38, 52, 14], F_20,513_= 8.20, cluster extent = 50, and in the right hemisphere, peak MNI coordinates [42, 52,16], F_20,513_= 10.42, cluster extent = 75 voxels.

Additionally, effects of demonstrative processing were also observed in the right frontal eye field, peak MNI coordinates [32, 6, 64], F_20,513_= 28.04, cluster extent = 46 voxels.

**Figure 6.**
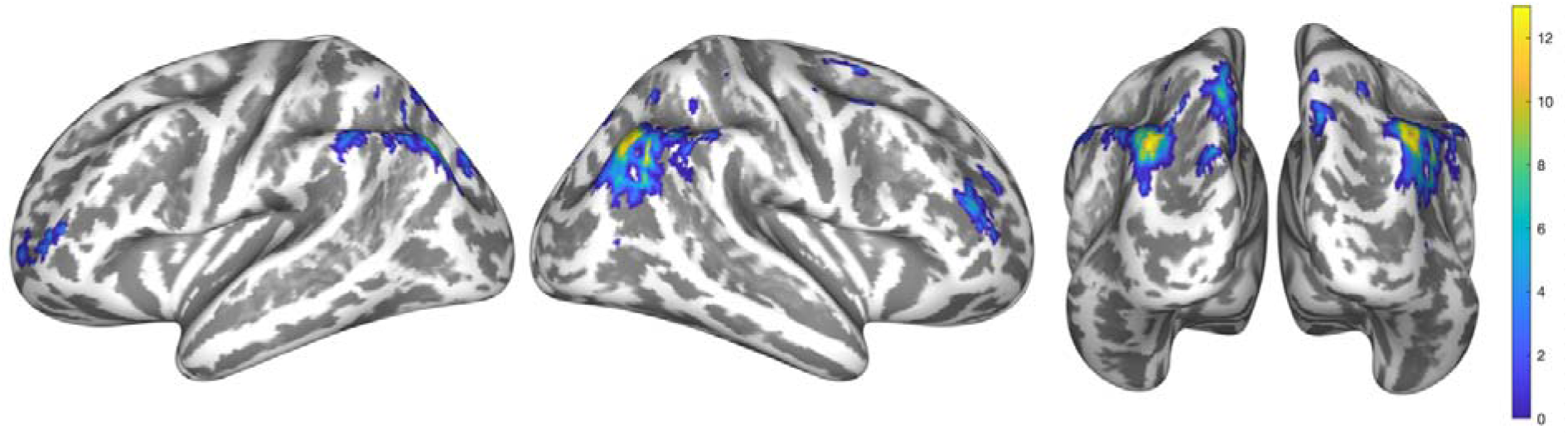
Brain regions responding to spatial demonstratives (both proximal and distal) across left and right channel. The analysis displays significant clusters in the inferior parietal cortices, in the medial superior parietal cortices, as well as in the middle frontal gyri, and right frontal eye field.

The time course of the response in parietal clusters is displayed in Figure 7. Response follows a slower time course than the auditory cortices, with peaks around 6 seconds after stimulus onset.

**Figure 7.**
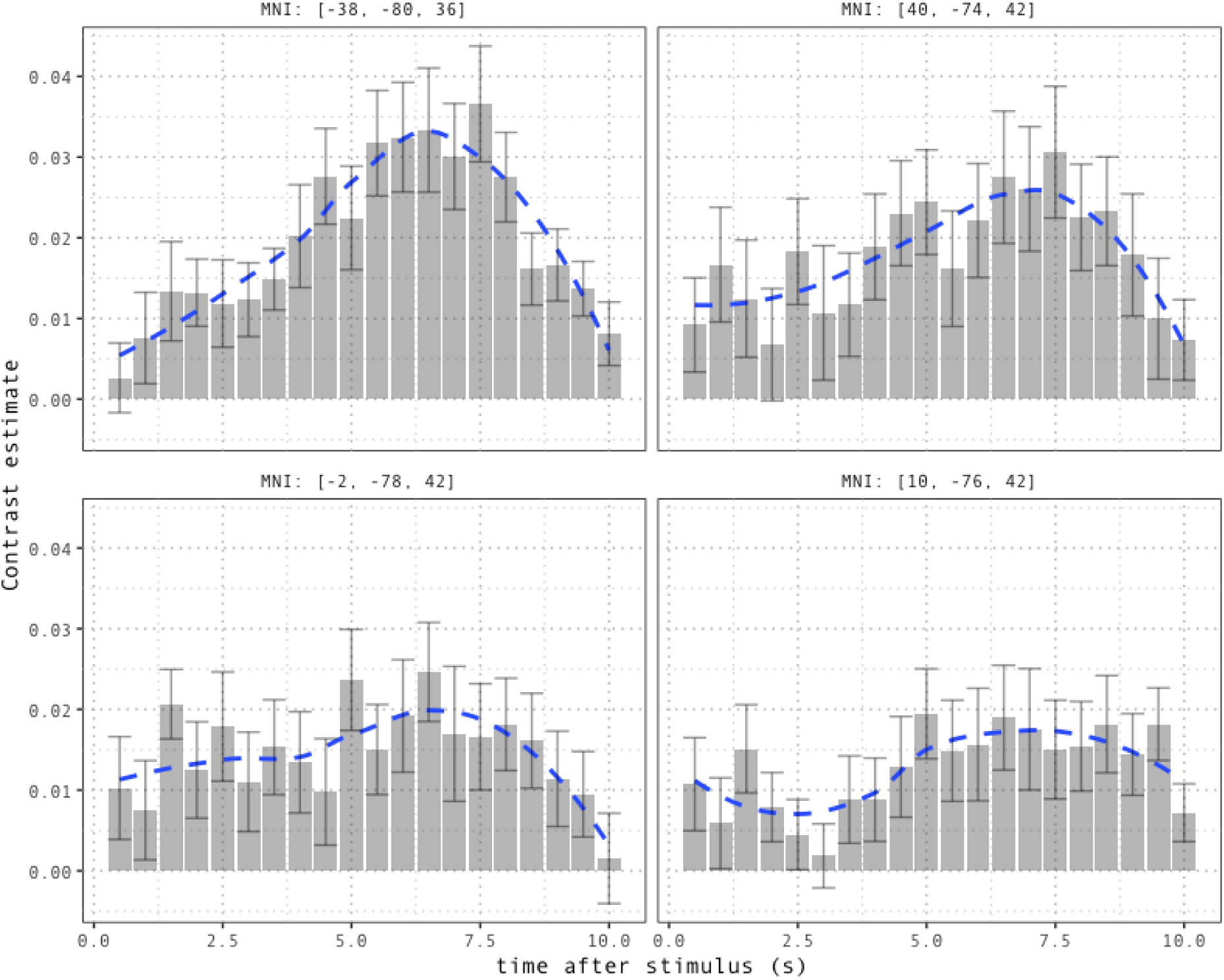
Time course of the response in peak voxels in the left angular gyrus (top left), right angular gyrus (top right), left precuneus (bottom left), right precuneus (bottom right). Error bars indicate between-participant standard errors. The overlaid curve smooths the average time series using local regression.

##### 3.1.2.2 Proximal vs. distal demonstratives

All the regions detected in the previous analysis were used as an inclusive mask for a direct comparison of distal and proximal demonstratives, aimed at highlighting differences between neural underpinnings of different demonstrative forms.

A direct comparison of proximal and distal demonstratives did not detect any significant cluster at a threshold of p < 0.05, and a cluster threshold of 30 voxels.

As a post-hoc test, we lowered the cluster threshold to 5 voxels to explore whether differences between proximal and distal demonstratives might be encoded in smaller neuronal subpopulations within the regions of interest.

The analysis displayed higher activation for distal demonstratives in clusters with peaks in the left angular gyrus, MNI: [−42, −78, 34], F_20,513_= 7.84, cluster extent = 13 voxels, right angular gyrus, MNI: [40, −74, 42], F_20,513_= 6.26, cluster extent = 13 voxels, right frontal eye fields, (MNI: [38, 6, 60], F_20,513_= 10.45, cluster extent = 12 voxels), and right middle frontal gyrus (MNI: [42, 52, 16], F_20,513_= 8.34, cluster extent = 8 voxels).

These patterns might indicate that responses to proximal and distal demonstrative differ in intensity (with larger response for distal demonstratives) rather than in neural substrates. However, given the lenient threshold used for this exploratory contrast, the small effect size, and since linguistic context for proximal and distal demonstrative forms was not controlled for in the text, these results provide a pointer for future studies, rather than direct evidence for the nature of semantic representation supporting different demonstrative forms.

##### 3.1.2.3 Whole-brain time course of response to demonstratives

Summarizing spatial and temporal features of neural response to demonstrative expressions, Figure 8 and Figure 9 display whole-brain parameter maps for proximal and distal demonstratives over contiguous 500ms time bins after word onset.

**Figure 8.**
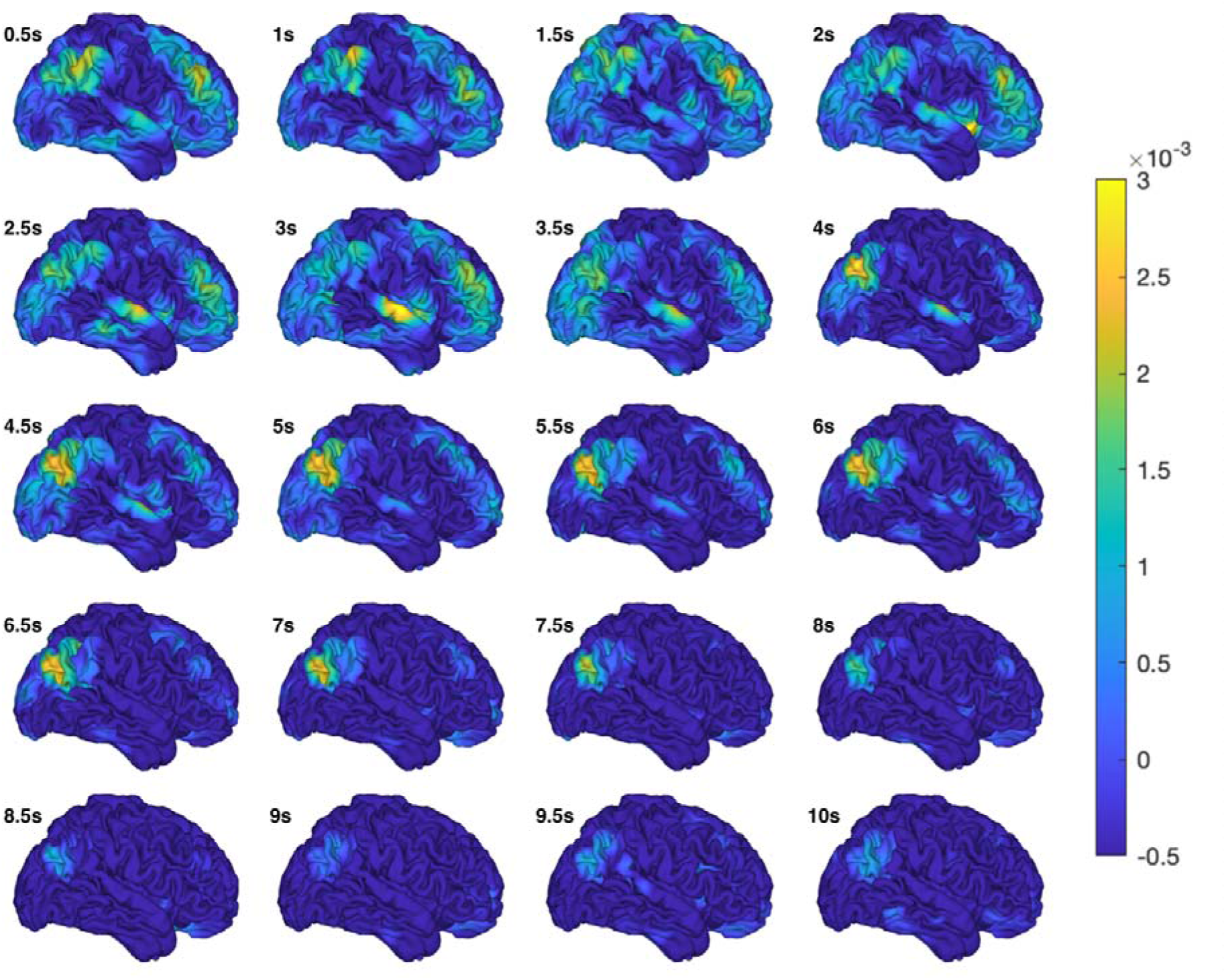
Parameter maps (averaged across participants) for *proximal* demonstratives over 10 seconds after stimulus onset, at 500ms intervals.

**Figure 9.**
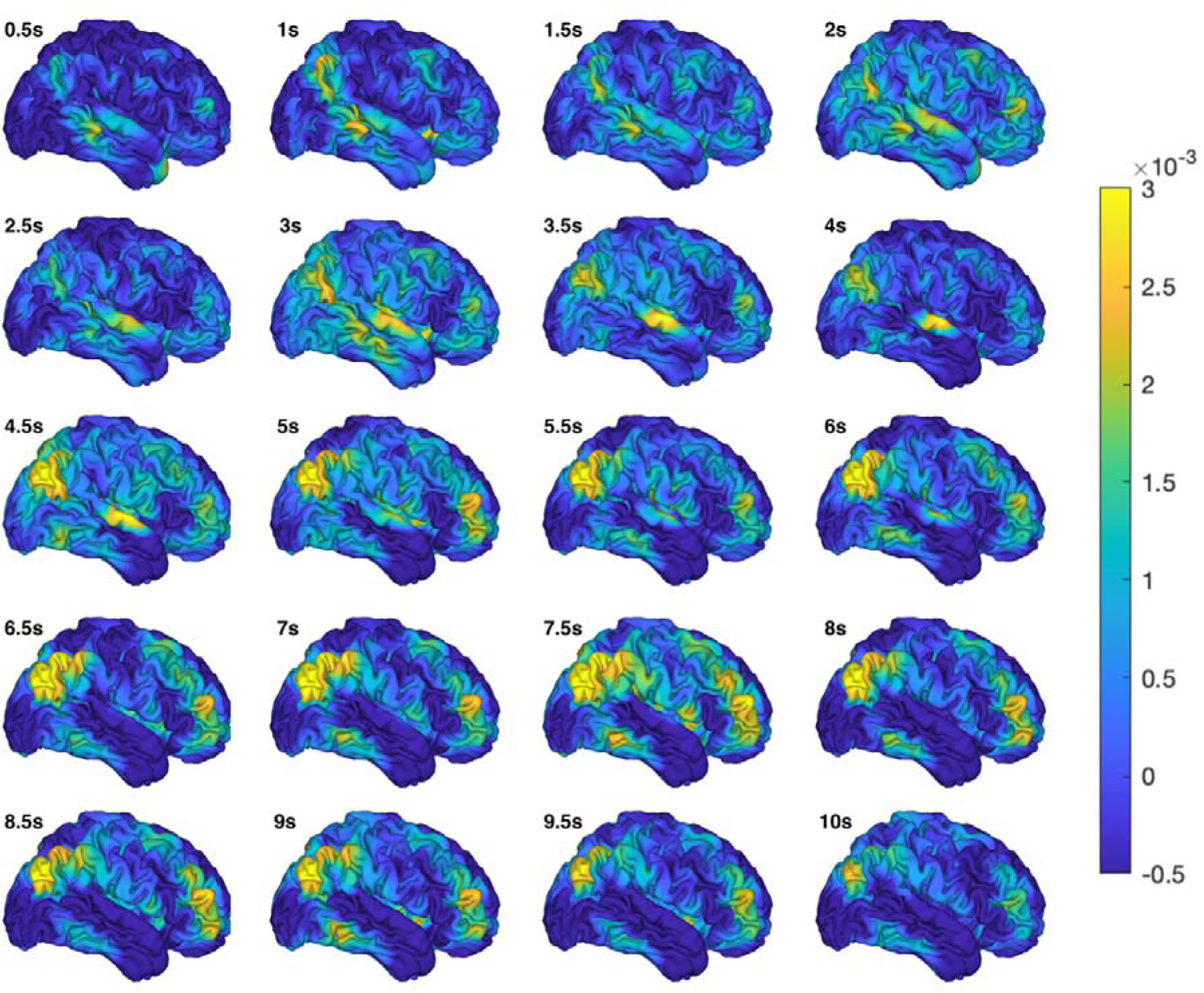
Parameter maps (averaged across participants) for *distal* demonstratives over 10 seconds after stimulus onset, at 500ms intervals.

Distal demonstratives exhibited more widespread and larger (although not significantly larger) responses than proximal demonstratives in all regions identified in the analysis. While the auditory cortices displayed an early and fast response, response in inferior parietal and medial occipital cortex peaks later in the case of proximal and distal demonstratives, with more sustained activation for distal demonstratives. Response in the frontal clusters showed higher-frequency fluctuations, with an early response for proximal demonstratives and multiple waves of activation for distal demonstratives.

##### 3.1.2.4 Interaction between demonstrative type and sound source

To identify whether any regions respond to the specific spatial location denoted by demonstratives, rather than to specific demonstrative forms, we tested for interactions between demonstrative form (proximal vs. distal) and sound source (left vs. right). As in the contrast between proximal and distal demonstratives, we constrained the analysis to those voxels that significantly responded to the occurrence of spatial demonstratives.

The rationale behind the test is that, if any areas respond more strongly to locations to the left of the participant, they would exhibit a positive response to both: a) occurrences of proximal demonstratives in the left channel; b) occurrences of distal demonstratives in the right channel, i.e. to instances of *here* or *this* uttered by the character located to the left of the participant, and instances of *there* or *that* uttered by the character located to the right of the participants. The opposite patterns would be observed for regions preferentially responding to locations in the right hemifield.

This contrast detected no significant voxels at a significance threshold of p(FWE-corrected) < .05 and a spatial extent threshold of 30 voxels, nor any clusters were detected when lowering the cluster threshold to 5 voxels.

#### 3.1.3 Wh-words

The occurrence of *where* in the text significantly modulated activity in clusters with peaks in the left angular gyrus, MNI: [−52, −62, 38], F_20,513_= 21.90, cluster extent = 93 voxels, and in the right angular gyrus, MNI: [44, −68, 42], F_20,513_= 10.60, cluster extent = 37 voxels. These clusters largely overlap with the inferior parietal clusters responding to the occurrence of spatial demonstratives. No clusters were detected when testing for effects of *what* and *who*.

### 3.2 Multivariate similarity analysis

#### 3.2.1 Whole-brain similarity between demonstratives and wh-words

Figure 10 displays between-participant averages of whole-brain topographical similarity (whole-brain correlations) between demonstratives and wh-word at each time point after stimulus onset.

**Figure 10.**
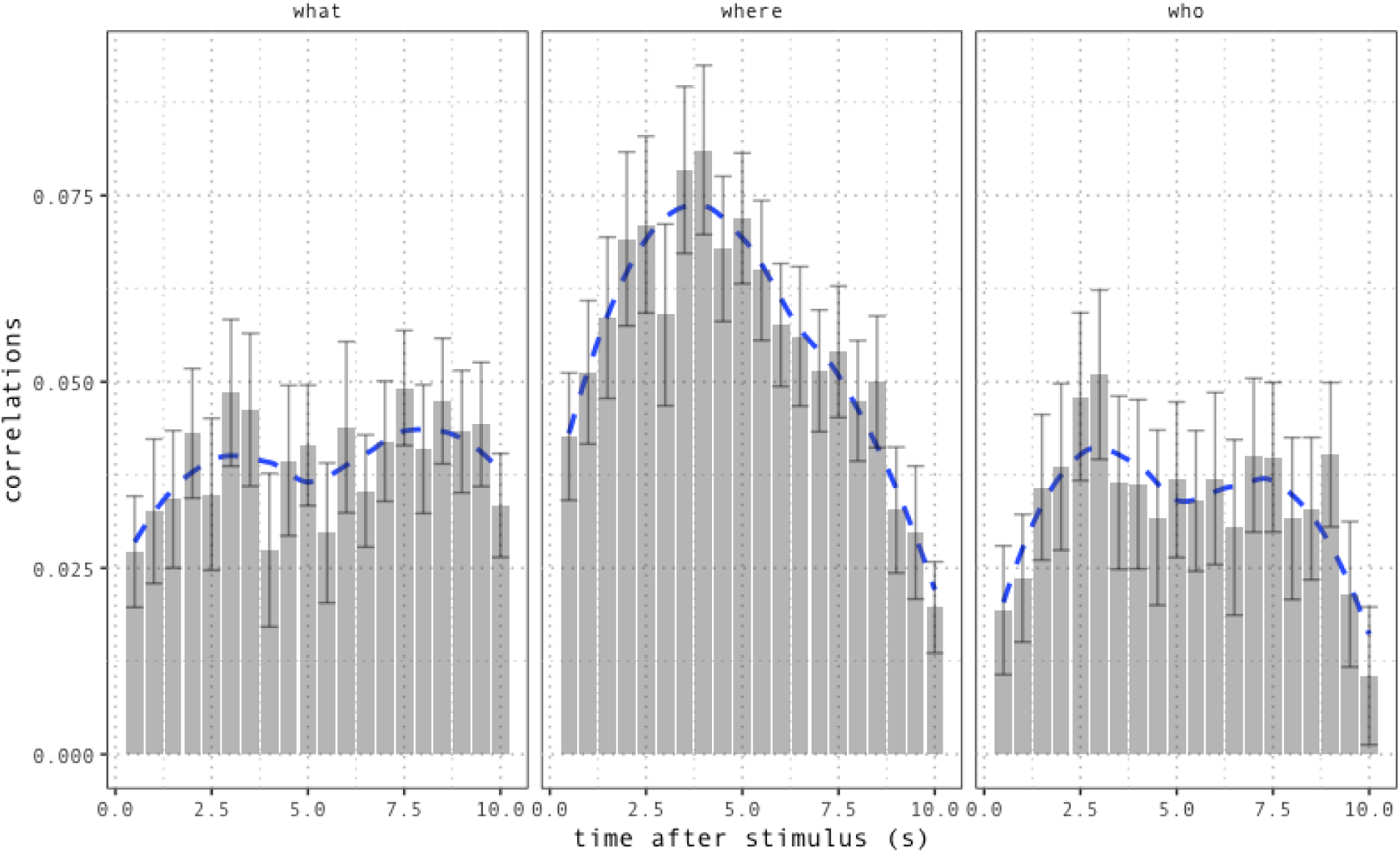
Whole-brain correlations between demonstratives and wh-words over time (500ms bins, over 10 seconds after stimulus onset). Bars denote averages across subjects and demonstrative type at each time point. The overlaid curve smooths the average time series using local regression. Error bars indicate standard error across participants. Correlations are on average higher for *where*, and their time course suggests similar BOLD response patterns for *where* and spatial demonstratives.

We extracted three summary metrics for the correlation time series. For each participant and each demonstrative/wh-word pair, we computed the area under the curve (AUC) defined by the correlation time series, as well as mean and maximum correlation over the 10 seconds span.

We used these measures to test for differences between wh-words in their overall topographical similarity with demonstratives using mixed-effects linear regressions. We fitted three models with the same fixed and random effects structure, and with AUC, mean correlation and maximum correlation as continuous outcome variables. In all models, the fixed effects structure included a categorical regressor coding for wh-word with *where* as reference level, while the random effects structure included an intercept for each subject and a random slope for the effect of wh-word.

In all models, similarity was higher for *where* compared to both *what* and *who*. AUC values were significantly lower for *what* compared to *where*, β = −0.16, se = 0.06, t(68.11) = −2.5, p < .05, and for *who* compared to *where*, β = −0.21, se = 0.07, t(27.41) = −3.07, p < .01. Post-hoc contrasts displayed no significant difference between *what* and *who*. Analogous patterns were observed using mean and maximum correlation as outcome variables (see Appendix C).

#### 3.2.2 Local similarity patterns

To zoom in on local topographical similarities and identify whether specific regions are driving the global similarity pattern observed above, we computed Pearson’s correlations between demonstratives and wh-words for 60 brain regions extracted from the AAL2 atlas, covering all regions within the frontal, temporal, parietal and occipital lobes. This yielded 28 (subjects) x 20 (time points) x 6 (word combinations) x 60 (regions of interest) similarity values. Here, correlation values represent topographical similarity between words within each of the regions.

Figure 11 provides an overview of correlations between neural representations of demonstratives and wh-words at each time point and for each brain region.

**Figure 11.**
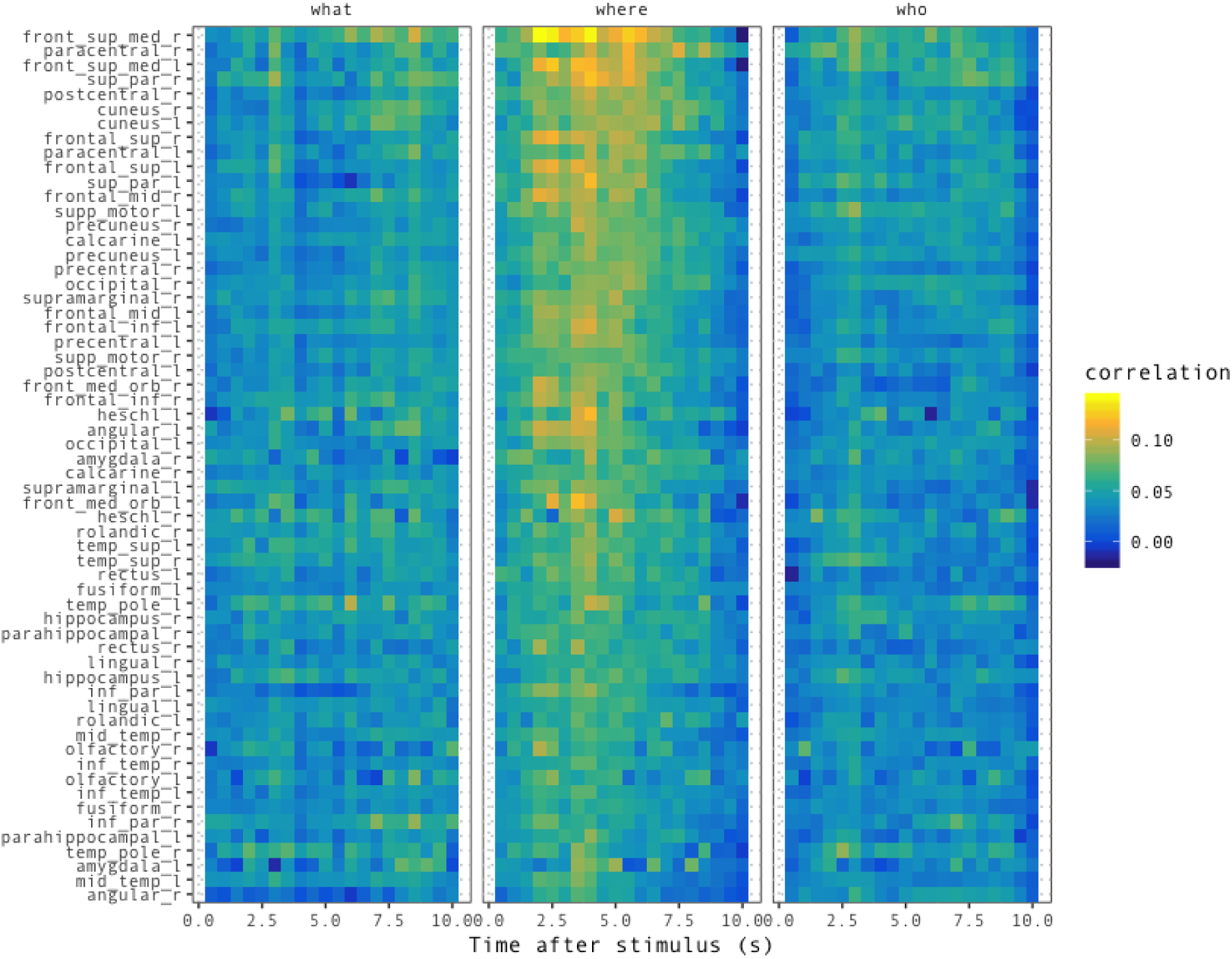
Local topographical similarity between demonstratives and wh-words over 10 seconds post stimulus onset in 60 brain regions extracted from the AAL2 atlas. Regions on the y-axis are sorted by ascending AUC for similarity between demonstratives and *where*.

The patterns in the figure suggest that correlations were lower for *what* and *who* compared to *where* across most regions, indicating that differences in topographical similarity at the whole-brain level reflect a widespread tendency rather than being uniquely driven by a small subset of regions.

Within this overall pattern, however, regions exhibit gradient variability. A group of frontal and parietal regions, located at the top of the graph (see Figure 11), displays markedly higher similarity with *where*, as well as a time course suggestive of analogous BOLD response patterns for demonstratives and *where*. These regions, bilaterally distributed and extending beyond the language network, largely overlap with the dorsal processing stream responsible for non-linguistic spatial perception, and they might constitute a network of neural resources for spatial cognition shared across the linguistic and non-linguistic domain.

**Figure 12.**
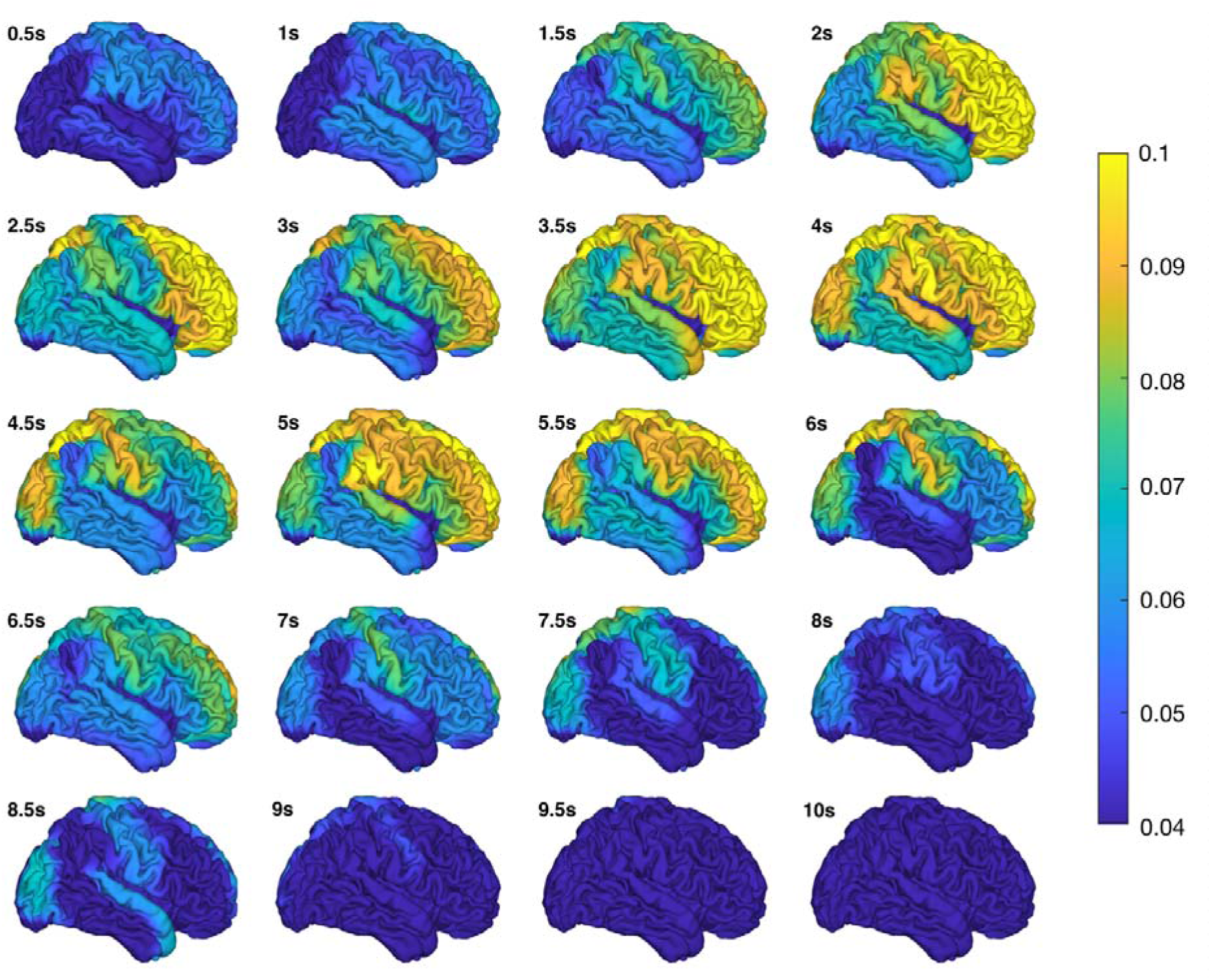
Correlation between demonstratives and *where* by region of interest over time. Colours code for average correlation values across subjects. Areas along the dorsal stream exhibit higher correlation values, with similarity evolving at a time course plausible for hemodynamic response, suggesting that the dorsal stream might constitute a network of resources for spatial processing shared across the linguistic and non-linguistic domain.

## 4. Discussion

Spatial demonstratives are powerful linguistic tools used to manipulate and share attention, and rely heavily on the synergy between language, perception, and spatial cognition. In this experiment, we investigated how this intertwining of linguistic and extra-linguistic cognition is implemented in the brain. This interplay is pivotal to language comprehension in general, but especially prominent in demonstratives. As predicted, we observed that spatial demonstratives engage a network of frontoparietal areas previously implicated in the construction, maintenance, and navigation of visuospatial representations. Additional analyses suggested that dorsal visuospatial pathways might be generally implicated in the processing of linguistic spatial expressions.

### 4.1 Integrating input, space and context in the posterior parietal cortex

Consistent with our predictions, demonstratives elicited bilateral responses in the supramarginal gyri, the posterior part of the angular gyrus, extending towards the middle occipital gyrus, as well as in medial superior parietal clusters with peaks in the precuneus. Crucially, all these regions are part of dorso-parietal visuospatial pathways not specific to linguistic processing (Kravitz, Saleem, Baker, & Mishkin, 2011).

The supramarginal gyrus is part of the temporo-parietal junction, responsible for interfacing the auditory cortex with parietal and frontal systems (Scott & Johnsrude, 2003). It is anatomically connected to the angular gyrus (Lee et al., 2007), a heteromodal association area (Bonner, Peelle, Cook, & Grossman, 2013; A. R. Damasio, 1989; Rademacher, Galaburda, Kennedy, Filipek, & Caviness Jr, 1992) implicated in a variety of processes requiring the integration of (task-relevant) information into coherent wholes (Seghier, 2013).

Integrating novel incoming information with previously constructed spatial and semantic contexts is crucial for spatial demonstratives. To decode the intended location, the coarse distance cues encoded by the semantics of specific forms (*here* = *near* vs. *there = far*) need to be integrated with knowledge on the position of the speaker within the previously constructed spatial scene, as well as with context-driven expectations on the intended referent. In this process, the angular gyri are supported by co-activated parietal clusters. Representations of spatial scenes are maintained in working memory and updated by the precuneus, which is directly connected to the angular gyrus via the occipitofrontal fascicle (Makris et al., 2007) and has previously been implicated in spatial working memory for both vision and language (Wallentin, Weed, et al., 2008; Zaehle et al., 2007).

### 4.2 Attentional orienting towards intended location: frontal clusters

Demonstratives bear a close link to attentional reorienting, as they are used to directly trigger attentional shifts towards relevant locations. Congruent with this, we found increased activation in the anterior part of the middle frontal gyrus bilaterally (BA10 and BA46) and in the right frontal eye fields. These areas belong to attentional networks responsible for controlling visually and memory-guided saccades also in absence of overt attentional reorienting (Corbetta et al., 1998; Fox, Corbetta, Snyder, Vincent, & Raichle, 2006). The frontal eye fields have previously been implicated in shifts in reference frames for the processing of linguistic spatial relations (Wallentin, 2012), a process relevant in decoding the referent of spatial demonstratives. Spatial demonstratives provide distance cues on the location of the intended referent relative to the speaker (or the dyad), thus requiring a transition from a default egocentric encoding of the scene to an allocentric frame with the speaker’s position as centre. However, significant activation in the frontal eye fields was only found in the right hemisphere, an asymmetry which does not directly resonate with previous studies and calls for further investigation. Other complementary factors might explain the effect of demonstratives on the FEF. FEF activation might also be triggered by participants performing actual eye movements in response to demonstratives. While this interpretation is compatible with the function of the frontal eye fields, it cannot be tested empirically in this context as no eye-tracking data were collected during the experiment.

### 4.3 No spatial segregation between proximal and distal forms

Contrary to our predictions, we did not find any evidence for spatially distinct substrates supporting processing of proximal and distal demonstratives, contrary to the hypothesis that areas coding for object reachability would be differentially recruited by the two demonstrative forms. This lack of spatial segregation might be explained by different factors. While reachability might be driving speakers’ choices in production, explicit encoding of reachability might not be necessary for comprehension. Scanning the visual scene on the basis of allocentric distance indications (near vs. far from the speaker), as well as the aid of context-driven expectations, might be sufficient to identify the intended referent.

A further explanation for the lack of segregation between demonstrative forms might be the absence of a clear-cut partition between neural resources coding for reachable and non-reachable locations. Solid evidence for such segregation has been found in non-human primates (Batista, Buneo, Snyder, & Andersen, 1999; Cohen & Andersen, 2002; Colby & Goldberg, 1999). In humans, behavioural patterns from visuo-tactile integration tasks are coherent with the existence of a similar architecture (di Pellegrino & Làdavas, 2015; Làdavas, 2002), and in line with neuropsychological evidence for double dissociations between peripersonal and extrapersonal neglect (Halligan et al., 2003; Halligan & Marshall, 1991; Ten Brink, Biesbroek, Oort, Visser-Meily, & Nijboer, 2019; Vuilleumier, Valenza, Mayer, Reverdin, & Landis, 1998). However, coherent evidence for a hard-wired segregation is yet to be found. A number of studies have attempted to identify areas exclusively associated to manual reach (Connolly et al., 2003; Gallivan et al., 2009), but object reachability is often confounded with purely visual parameters, such as distance from the subject and position of the target relative to the centre of fixation.

Nonetheless, we detected magnitude differences in response to proximal and distal forms in all areas responding to demonstratives. This finding might be explained by distal forms imposing heavier processing demands. Proximal expressions denote a location which is roughly equivalent to the position of the speaker. Shifting attentional focus towards the speaker might be enough to decode the intended referent. On the other hand, distal demonstratives provide more underspecified cues, compatible with any location which is *not* in the proximity of the speaker. In this case, reference resolution might require a more extensive search, and more heavily rely on the integration between spatial context and top-down expectations. Further experiments are needed to directly test this hypothesis.

Finally, we found no interactions between demonstrative forms and side of presentation, thus not supporting the hypothesis of areas of interest displaying preferences for the contralateral hemispace. Variability in the spatial configuration of the imagined scene across participants and the spatial underspecification of locations denoted by demonstrative forms (especially distal forms) in absence of an external visual stimulus might explain the lack of such an effect.

### 4.4 Spatial language and the dorsal stream

In our analysis, we showed that global topographical similarity between demonstratives and wh-words priming processing of spatial (*where*) content is higher than similarity with non-spatial (*what* and *who*) wh-words. This pattern is driven by frontal and parietal areas belonging to the dorsal stream and related pathways. We interpret this as suggestive of a functional role for dorsal (*where* or *how*, see Goodale & Milner, 1992) pathway(s) in the processing of linguistic expressions describing spatial relations, as opposed to ventral structures (the *what* stream) supporting semantic and conceptual processing.

The involvement (and functional segregation) of the *where* and *what* pathways in language comprehension has previously been hypothesized on a theoretical basis (Landau & Jackendoff, 1993, 2013), and has been indirectly supported by empirical evidence on the overlap between neural substrates for linguistic and visual spatial processing (Wallentin, Weed, et al., 2008). However, our study is the first to provide direct evidence of a general involvement of the dorsal stream in spatial language, and it paves the way for further research.

The dorsal stream might not be recruited exclusively by spatial expressions, but rather exhibit a preference for words heavily relying on contextual integration. Moreover, some studies have suggested a tripartite organization of the dorsal stream into pathways encoding spatial information for manual action (parieto-premotor pathways), attentional orienting (parietal-prefrontal pathway), and spatial navigation (parieto-medial-temporal pathway) (Kravitz et al., 2011). This tripartite distinction might also be reflected in a further functional specialization of dorsal pathways for different types of linguistic spatial reference frames (e.g. allocentric or landmark-based vs. egocentric reference, categorical vs. coordinate-based encoding).

Finally, direct involvement of the dorsal stream in spatial language might bear a crucial indication on the nature of the neurobiological substrates of language processing in line with distributed accounts (Barsalou, Simmons, Barbey, & Wilson, 2003; Desai, Choi, Lai, & Henderson, 2016; Fernandino et al., 2015; Huth, Nishimoto, Vu, & Gallant, 2012; Pulvermüller, 2005). Rather than relying on a specialized circuitry, language processing seems to engage a flexible and non-segregated architecture, where neural structures supporting perceptual, attentional and higher-level cognitive tasks are dynamically recruited and mutually interfaced in a context-dependent fashion.

## 5. Conclusions

We conducted a naturalistic fast fMRI experiment to investigate the neural correlates of spatial demonstratives. Our findings suggest that processing spatial demonstratives recruits dorsal parieto-frontal areas previously implicated in extra-linguistic visuospatial cognition and attentional orienting. Additionally, we provide evidence that dorsal “where” pathways might be generally involved in the processing of linguistic spatial expressions, as opposed to ventral pathways encoding object semantics. More generally, these results suggest that language processing might rely on a distributed and non-segregated architecture, recruiting neural resources for attention, perception and extra-linguistic aspects of cognition in a dynamic and context-dependent fashion.

## Funding

This experiment was funded by the DCOMM project (European Union Horizon 2020 research and innovation programme, Marie Skłodowska-Curie Actions; grant agreement 676063).

## Acknowledgements

We would like to thank the Center for Magnetic Resonance Research, University of Minnesota for providing access to the multiband-EPI sequence.

## Competing interests

The authors declare no competing interests.

## Appendix A

The 60 regions of interest included in the similarity analysis correspond to all sub-regions of the frontal, temporal, parietal and occipital lobes in AAL2 (see Table 2 in Rolls et al., 2015). The opercular, triangular and orbital part of the inferior frontal gyrus were collapsed into a single region of interest. Homologue regions in left and right hemisphere were kept as separate ROIs.

## Appendix B

In addition to the analyses targeting the neural correlates of spatial demonstratives, we reported further analyses aimed at ensuring the quality of the images yielded by our acquisition sequence and exploring lateralization patterns for auditory response to the speech stimulus.

We fitted sound envelopes from the left and right auditory channel to the EPI time series expecting to detect robust effects of the low-level profile of the signal in the auditory cortices.

As predicted, auditory cortices responded bilaterally to monaural input, but, for both ears, response in the contralateral hemisphere was larger than response in the ipsilateral, which is consistent with previous studies (Hirano et al., 1997; Jäncke et al., 2002; Schönwiesner et al., 2006; Stefanatos et al., 2008; Suzuki et al., 2002).

Interestingly, when directly comparing the effects of the left and the right envelope, we observed asymmetries across the two hemispheres. Response in the left auditory cortex was significantly larger for input from the contralateral ear than from the ipsilateral. However, no such difference was observed for the right auditory cortex, where the magnitude of the response was comparable across ears. A similar lateralization pattern has been previously reported for non-speech stimuli (Schönwiesner et al., 2006), and it might be compatible with different interpretations. Input from the right ear seems to elicit larger and more widespread response in both auditory cortices.

This levels out responses to right and left ear in the right hemisphere, while preserving an advantage for the right ear in the left auditory cortex. Larger overall responsiveness to right-ear input might explain the widely-attested behavioural advantage for right-ear input observed in dichotic listening (Hugdahl & Westerhausen, 2016; Kimura, 1967). Functional specialization of right and left auditory cortices for fine spectral and temporal features respectively might also be compatible with our results (Zatorre, Belin, & Penhune, 2002). The analytic envelope of the sound preserves spectral modulations of the signal while filtering out its fine temporal structure. Specialization for spectral features might therefore explain why the magnitude of response to sound envelopes in the right AC remains constant regardless of spatial origin of the sound.

Finally, the temporal profile of contrast estimates from the FIR analysis showed that response in the primary AC peaked between 3 and 4 seconds after stimulus onset, earlier than reported in previous literature (Hall et al., 2000). This pattern is compatible with previous studies showing faster response in primary sensory areas for sustained and rapidly varying input (Lewis et al., 2016). Further analyses are needed to achieve a reliable characterization of time course of the signal under naturalistic conditions.

## Appendix C

Supplementary analyses on whole-brain similarity between demonstratives and wh-words using mean and maximum correlation as outcome variables displayed results analogous to those obtained using AUC as outcome. Mean correlations were significantly lower for *what* compared to *where*, β = −0.02, se = 0.007, t(69.74) = −2.43, p < .05, and for *who* compared to *where*, β = −0.02, se = 0.007, t(27.27) = −3.11, p < .01. The same effects were detected using maximum correlation as outcome, with significantly lower correlations for *what* than for *where*, β = −0.02, se = 0.008, t(36.22) = −2.77, p < .01, and for *who* than for *where*, β = −0.03, se = 0.009, t(31) = −3.54, p < .01. Post-hoc contrasts displayed no significant difference for *what* and *who* on any of the outcome measures.

## Notes

#### Summary of Updates

Addition of details concerning previous studies on demonstratives, clarification on aspects of the methodology and the stimuli, replacement of all figures with figures using colorblind-friendly colormap, backgrounding of the discussion on sound envelope results by moving the relevant paragraph to the appendix.

https://osf.io/j9fm5/

## References

Andersen, R. A., Andersen, K. N., Hwang, E. J., & Hauschild, M. (2014). Optic ataxia: From Balint’s syndrome to the parietal reach region. Neuron, 81(5), 967–983.

Barsalou, L. W., Simmons, W. K., Barbey, A. K., & Wilson, C. D. (2003). Grounding conceptual knowledge in modality-specific systems. Trends in Cognitive Sciences, 7(2), 84–91.

Batista, A. P., Buneo, C. A., Snyder, L. H., & Andersen, R. A. (1999). Reach plans in eye-centered coordinates. Science, 285(5425), 257–260.

Binder, J. R., Desai, R. H., Graves, W. W., & Conant, L. L. (2009). Where is the semantic system? A critical review and meta-analysis of 120 functional neuroimaging studies. Cerebral Cortex, 19(12), 2767–2796.

Bollmann, S., Puckett, A. M., Cunnington, R., & Barth, M. (2018). Serial correlations in single-subject fMRI with sub-second TR. NeuroImage, 166, 152–166.

Bonfiglioli, C., Finocchiaro, C., Gesierich, B., Rositani, F., & Vescovi, M. (2009). A kinematic approach to the conceptual representations of this and that. Cognition, 111(2), 270–274.

Bonner, M. F., Peelle, J. E., Cook, P. A., & Grossman, M. (2013). Heteromodal conceptual processing in the angular gyrus. Neuroimage, 71, 175–186.

Caldano, M., & Coventry, K. R. (in press). Spatial demonstratives and perceptual space. To reach or not to reach in the sagittal and lateral planes? Cognition.

Chang, C., Cunningham, J. P., & Glover, G. H. (2009). Influence of heart rate on the BOLD signal: The cardiac response function. Neuroimage, 44(3), 857–869.

Clark, E. V., & Sengul, C. J. (1978). Strategies in the acquisition of deixis. Journal of Child Language, 5(3), 457–475.

Cohen, Y. E., & Andersen, R. A. (2002). A common reference frame for movement plans in the posterior parietal cortex. Nature Reviews Neuroscience, 3(7), 553.

Colby, C. L., & Goldberg, M. E. (1999). Space and attention in parietal cortex. Annual Review of Neuroscience, 22(1), 319–349.

Connolly, J. D., Andersen, R. A., & Goodale, M. A. (2003). FMRI evidence for a’parietal reach region’in the human brain. Experimental Brain Research, 153(2), 140–145.

Cooperrider, K. (2016). The co-organization of demonstratives and pointing gestures. Discourse Processes, 53(8), 632–656.

Corbetta, M., Akbudak, E., Conturo, T. E., Snyder, A. Z., Ollinger, J. M., Drury, H. A., … Van Essen, D. C. (1998). A common network of functional areas for attention and eye movements. Neuron, 21(4), 761–773.

Coventry, K. R., Griffiths, D., & Hamilton, C. J. (2014). Spatial demonstratives and perceptual space: Describing and remembering object location. Cognitive Psychology, 69, 46–70.

Coventry, K. R., Valdés, B., Castillo, A., & Guijarro-Fuentes, P. (2008). Language within your reach: Near–far perceptual space and spatial demonstratives. Cognition, 108(3), 889–895.

Damasio, A. R. (1989). The brain binds entities and events by multiregional activation from convergence zones. Neural Computation, 1(1), 123–132.

Damasio, H., Grabowski, T. J., Tranel, D., Ponto, L. L., Hichwa, R. D., & Damasio, A. R. (2001). Neural correlates of naming actions and of naming spatial relations. Neuroimage, 13(6), 1053–1064.

Desai, R. H., Choi, W., Lai, V. T., & Henderson, J. M. (2016). Toward semantics in the wild: Activation to manipulable nouns in naturalistic reading. Journal of Neuroscience, 36(14), 4050–4055.

di Pellegrino, G., & Làdavas, E. (2015). Peripersonal space in the brain. Neuropsychologia, 66, 126–133.

Diessel, H. (1999). Demonstratives: Form, function and grammaticalization (Vol. 42). John Benjamins Publishing.

Diessel, H. (2006). Demonstratives, joint attention, and the emergence of grammar. Cognitive Linguistics, 17(4), 463–489.

Diessel, H. (2013). Where does language come from? Some reflections on the role of deictic gesture and demonstratives in the evolution of language. Language and Cognition, 5(2–3), 239–249.

Diessel, H. (2014). Demonstratives, frames of reference, and semantic universals of space. Language and Linguistics Compass, 8(3), 116–132.

Fernandino, L., Binder, J. R., Desai, R. H., Pendl, S. L., Humphries, C. J., Gross, W. L., … Seidenberg, M. S. (2015). Concept representation reflects multimodal abstraction: A framework for embodied semantics. Cerebral Cortex, 26(5), 2018–2034.

Fox, M. D., Corbetta, M., Snyder, A. Z., Vincent, J. L., & Raichle, M. E. (2006). Spontaneous neuronal activity distinguishes human dorsal and ventral attention systems. Proceedings of the National Academy of Sciences, 103(26), 10046–10051.

Gallivan, J. P., Cavina-Pratesi, C., & Culham, J. C. (2009). Is that within reach? FMRI reveals that the human superior parieto-occipital cortex encodes objects reachable by the hand. Journal of Neuroscience, 29(14), 4381–4391.

García, J. O. P., Ehlers, K. R., & Tylén, K. (2017). Bodily constraints contributing to multimodal referentiality in humans: The contribution of a de-pigmented sclera to proto-declaratives. Language & Communication, 54, 73–81.

Glover, G. H., Li, T.-Q., & Ress, D. (2000). Image□based method for retrospective correction of physiological motion effects in fMRI: RETROICOR. Magnetic Resonance in Medicine: An Official Journal of the International Society for Magnetic Resonance in Medicine, 44(1), 162–167.

Goodale, M. A., & Milner, A. D. (1992). Separate visual pathways for perception and action. Trends in Neurosciences, 15(1), 20–25.

Grivaz, P., Blanke, O., & Serino, A. (2017). Common and distinct brain regions processing multisensory bodily signals for peripersonal space and body ownership. Neuroimage, 147, 602–618.

Gudde, H. B., Coventry, K. R., & Engelhardt, P. E. (2016). Language and memory for object location. Cognition, 153, 99–107.

Hall, D. A., Summerfield, A. Q., Gonçalves, M. S., Foster, J. R., Palmer, A. R., & Bowtell, R. W. (2000). Time□course of the auditory BOLD response to scanner noise. Magnetic Resonance in Medicine: An Official Journal of the International Society for Magnetic Resonance in Medicine, 43(4), 601–606.

Halligan, P. W., Fink, G. R., Marshall, J. C., & Vallar, G. (2003). Spatial cognition: Evidence from visual neglect. Trends in Cognitive Sciences, 7(3), 125–133.

Halligan, P. W., & Marshall, J. C. (1991). Left neglect for near but not far space in man. Nature, 350(6318), 498.

Hassabis, D., & Maguire, E. A. (2009). The construction system of the brain. Philosophical Transactions of the Royal Society B: Biological Sciences, 364(1521), 1263–1271.

Henson, R. N. A. (2003). Analysis of fMRI time series: Linear time-invariant models, event-related fMRI and optimal experimental design. Elsevier.

Hugdahl, K., & Westerhausen, R. (2016). Speech processing asymmetry revealed by dichotic listening and functional brain imaging. Neuropsychologia, 93, 466–481.

Huth, A. G., Nishimoto, S., Vu, A. T., & Gallant, J. L. (2012). A continuous semantic space describes the representation of thousands of object and action categories across the human brain. Neuron, 76(6), 1210–1224.

Jäncke, L., Wüstenberg, T., Schulze, K., & Heinze, H. J. (2002). Asymmetric hemodynamic responses of the human auditory cortex to monaural and binaural stimulation. Hearing Research, 170(1–2), 166–178.

Kasper, L., Bollmann, S., Diaconescu, A. O., Hutton, C., Heinzle, J., Iglesias, S., … Pruessmann, K. P. (2017). The PhysIO toolbox for modeling physiological noise in fMRI data. Journal of Neuroscience Methods, 276, 56–72.

Kemmerer, D. (1999). “Near” and “far” in language and perception. Cognition, 73(1), 35–63.

Kemmerer, D. (2006). The semantics of space: Integrating linguistic typology and cognitive neuroscience. Neuropsychologia, 44(9), 1607–1621.

Kimura, D. (1967). Functional asymmetry of the brain in dichotic listening. Cortex, 3(2), 163–178.

Kravitz, D. J., Saleem, K. S., Baker, C. I., & Mishkin, M. (2011). A new neural framework for visuospatial processing. Nature Reviews Neuroscience, 12(4), 217.

Làdavas, E. (2002). Functional and dynamic properties of visual peripersonal space. Trends in Cognitive Sciences, 6(1), 17–22.

Landau, B., & Jackendoff, R. (1993). Whence and whither in spatial language and spatial cognition? Behavioral and Brain Sciences, 16(2), 255–265.

Landau, B., & Jackendoff, R. (2013). Spatial language and spatial cognition. In Bridges between psychology and linguistics (pp. 157–182). Psychology Press.

Lee, H., Devlin, J. T., Shakeshaft, C., Stewart, L. H., Brennan, A., Glensman, J., … Frackowiak, R. S. (2007). Anatomical traces of vocabulary acquisition in the adolescent brain. Journal of Neuroscience, 27(5), 1184–1189.

Leech, G., & Rayson, P. (2014). Word frequencies in written and spoken English: Based on the British National Corpus. Routledge.

Levinson, S. C. (1983). Pragmatics. Cambridge textbooks in linguistics. Cambridge/New York.

Lewis, L. D., Setsompop, K., Rosen, B. R., & Polimeni, J. R. (2016). Fast fMRI can detect oscillatory neural activity in humans. Proceedings of the National Academy of Sciences, 113(43), E6679–E6685.

Lund, T. E., Madsen, K. H., Sidaros, K., Luo, W.-L., & Nichols, T. E. (2006). Non-white noise in fMRI: does modelling have an impact? Neuroimage, 29(1), 54–66.

Makris, N., Papadimitriou, G. M., Sorg, S., Kennedy, D. N., Caviness, V. S., & Pandya, D. N. (2007). The occipitofrontal fascicle in humans: A quantitative, in vivo, DT-MRI study. Neuroimage, 37(4), 1100–1111.

Mishkin, M., Ungerleider, L. G., & Macko, K. A. (1983). Object vision and spatial vision: Two cortical pathways. Trends in Neurosciences, 6, 414–417.

Noordzij, M. L., Neggers, S. F., Ramsey, N. F., & Postma, A. (2008). Neural correlates of locative prepositions. Neuropsychologia, 46(5), 1576–1580.

Peeters, D., Hagoort, P., & Özyürek, A. (2015). Electrophysiological evidence for the role of shared space in online comprehension of spatial demonstratives. Cognition, 136, 64–84.

Peirce, J. W. (2007). PsychoPy—psychophysics software in Python. Journal of Neuroscience Methods, 162(1–2), 8–13.

Pulvermüller, F. (2005). Brain mechanisms linking language and action. Nature Reviews Neuroscience, 6(7), 576.

Purdon, P. L., & Weisskoff, R. M. (1998). Effect of temporal autocorrelation due to physiological noise and stimulus paradigm on voxel□level false□positive rates in fMRI. Human Brain Mapping, 6(4), 239–249.

Rademacher, J., Galaburda, A. M., Kennedy, D. N., Filipek, P. A., & Caviness Jr, V. S. (1992). Human cerebral cortex: Localization, parcellation, and morphometry with magnetic resonance imaging. Journal of Cognitive Neuroscience, 4(4), 352–374.

Rocca, R., Tylén, K., & Wallentin, M. (2019). This shoe, that tiger: Semantic properties reflecting manual affordances of the referent modulate demonstrative use. PLOS ONE, 14(1), e0210333.

Rocca, R., Wallentin, M., Vesper, C., & Tylén, K. (2018). This and that back in context: Grounding demonstrative reference in manual and social affordances. Proceedings of The 40th Annual Meeting Of The Cognitive Science Society, Madison, Wisconsin.

Rocca, R., Wallentin, M., Vesper, C., & Tylén, K. (2019). This is for you: Social modulations of proximal vs. Distal space in collaborative interactions.

Rolls, E. T., Joliot, M., & Tzourio-Mazoyer, N. (2015). Implementation of a new parcellation of the orbitofrontal cortex in the automated anatomical labeling atlas. Neuroimage, 122, 1–5.

Sahib, A. K., Mathiak, K., Erb, M., Elshahabi, A., Klamer, S., Scheffler, K., … Ethofer, T. (2016). Effect of temporal resolution and serial autocorrelations in event□related functional MRI. LJ Magnetic Resonance in Medicine, 76(6), 1805–1813.

Schönwiesner, M., Krumbholz, K., Rübsamen, R., Fink, G. R., & von Cramon, D. Y. (2006). Hemispheric asymmetry for auditory processing in the human auditory brain stem, thalamus, and cortex. Cerebral Cortex, 17(2), 492–499.

Scott, S. K., & Johnsrude, I. S. (2003). The neuroanatomical and functional organization of speech perception. Trends in Neurosciences, 26(2), 100–107.

Seghier, M. L. (2013). The Angular Gyrus:Multiple Functions and Multiple Subdivisions. The Neuroscientist, 19(1), 43–61.

Setsompop, K., Cohen-Adad, J., Gagoski, B. A., Raij, T., Yendiki, A., Keil, B., … Wald, L. L. (2012). Improving diffusion MRI using simultaneous multi-slice echo planar imaging. Neuroimage, 63(1), 569–580.

Stefanatos, G. A., Joe, W. Q., Aguirre, G. K., Detre, J. A., & Wetmore, G. (2008). Activation of human auditory cortex during speech perception: Effects of monaural, binaural, and dichotic presentation. Neuropsychologia, 46(1), 301–315.

Stevens, J., & Zhang, Y. (2013). Relative distance and gaze in the use of entity-referring spatial demonstratives: An event-related potential study. Journal of Neurolinguistics, 26(1), 31–45.

Ten Brink, A. F., Biesbroek, J. M., Oort, Q., Visser-Meily, J. M., & Nijboer, T. C. (2019). Peripersonal and extrapersonal visuospatial neglect in different frames of reference: A brain lesion-symptom mapping study. Behavioural Brain Research, 356, 504–515.

Tomasello, M., Carpenter, M., Call, J., Behne, T., & Moll, H. (2005). In search of the uniquely human. Behavioral and Brain Sciences, 28(5), 721–727.

Tylén, K., Weed, E., Wallentin, M., Roepstorff, A., & Frith, C. D. (2010). Language as a tool for interacting minds. Mind & Language, 25(1), 3–29.

Tzourio-Mazoyer, N., Landeau, B., Papathanassiou, D., Crivello, F., Etard, O., Delcroix, N., … Joliot, M. (2002). Automated anatomical labeling of activations in SPM using a macroscopic anatomical parcellation of the MNI MRI single-subject brain. Neuroimage, 15(1), 273–289.

Vuilleumier, P., Valenza, N., Mayer, E., Reverdin, A., & Landis, T. (1998). Near and far visual space in unilateral neglect. Annals of Neurology, 43(3), 406–410.

Wallentin, M. (2009). Putative sex differences in verbal abilities and language cortex: A critical review. Brain and Language, 108(3), 175–183.

Wallentin, M. (2012). The role of the brain’s frontal eye fields in constructing frame of reference. Cognitive Processing, 13(1), 359–363.

Wallentin, M. (2018). Sex differences in post-stroke aphasia rates are caused by age. A meta-analysis and database query. PloS One, 13(12), e0209571.

Wallentin, M., Kristensen, L. B., Olsen, J. H., & Nielsen, A. H. (2011). Eye movement suppression interferes with construction of object-centered spatial reference frames in working memory. Brain and Cognition, 77(3), 432–437.

Wallentin, M., Roepstorff, A., & Burgess, N. (2008). Frontal eye fields involved in shifting frame of reference within working memory for scenes. Neuropsychologia, 46(2), 399–408.

Wallentin, M., Roepstorff, A., Glover, R., & Burgess, N. (2006). Parallel memory systems for talking about location and age in precuneus, caudate and Broca’s region. Neuroimage, 32(4), 1850–1864.

Wallentin, M., Weed, E., Østergaard, L., Mouridsen, K., & Roepstorff, A. (2008). Accessing the mental space—spatial working memory processes for language and vision overlap in precuneus. Human Brain Mapping, 29(5), 524–532.

Zaehle, T., Jordan, K., Wüstenberg, T., Baudewig, J., Dechent, P., & Mast, F. W. (2007). The neural basis of the egocentric and allocentric spatial frame of reference. Brain Research, 1137, 92–103.

Zatorre, R. J., Belin, P., & Penhune, V. B. (2002). Structure and function of auditory cortex: Music and speech. Trends in Cognitive Sciences, 6(1), 37–46.

